# Proteome-wide crosslinking mass spectrometry reveals novel components of essential complexes in *Toxoplasma*

**DOI:** 10.64898/2026.09.01.748362

**Authors:** Simon Butterworth, Ashley L. Gin, Shikha Shikha, Isadonna Tengganu, John Rush, Tom Duraisingh, Victoria Sodeinde, Leandro Lemgruber, Fabian Schulte, Ke Hu, Lilach Sheiner, Sergey Ovchinnikov, Sebastian Lourido

## Abstract

Protein–protein interactions underpin nearly all cellular processes, yet systematic definition of these networks remains limited outside a few model organisms. As a result, the architectures of essential complexes in many divergent lineages remain poorly characterized. Here we developed a high-coverage crosslinking mass spectrometry framework to map the proteome-wide interactome of the model apicomplexan parasite *Toxoplasma gondii*. From 29,624 crosslinked peptide pairs, we resolved a network of 2,859 protein–protein interactions that we integrated with structural modeling to resolve interaction interfaces. We identified and validated previously unrecognized components of essential protein complexes, including a structurally distinct ATP synthase subcomplex containing a highly divergent, apicomplexan-specific α subunit essential for parasite fitness. Beyond revealing unexpected diversification of core mitochondrial machinery, these findings provide a general strategy to define the molecular architecture of divergent organisms and represent a foundational resource for hypothesis generation, structural inference, and discovery of lineage-specific vulnerabilities in pathogen biology.

## INTRODUCTION

Most proteins function as components of oligomeric complexes^1,2^, which enable functional complexity and regulatory control that cannot be achieved by monomeric proteins. New biological functions can arise through modulation of individual subunits, addition of novel components, or replacement of existing ones within a complex. For poorly characterized proteins, interaction partners provide critical clues to function. Additionally, many diseases arise from aberrant protein–protein interactions due to mutation^3^, misfolding^4^, or interference by pathogen proteins^5,6^. Reliable identification of physiological protein–protein interactions is thus of central importance to understanding both normal cellular function and disease mechanisms.

Most technologies for mapping protein interactions are limited in one of two ways: targeted approaches such as co-immunoprecipitation and proximity labeling require genetic manipulation or custom antibodies—constraints that are particularly prohibitive in non-model organisms— while high-throughput alternatives like yeast two-hybrid sacrifice native cellular context entirely^7^. By contrast, crosslinking mass spectrometry (XL-MS) enables high-throughput identification of protein–protein interactions directly from native biological samples without the need for genetic engineering, capturing interactions in the cellular context in which they occur^8^. In XL-MS, a protein or cellular sample is treated with a bifunctional chemical crosslinker that forms covalent bonds between amino acids in close spatial proximity, capturing contacts within single proteins and between subunits of protein complexes. These covalent linkages are preserved through proteolytic digestion and can be identified by tandem mass spectrometry as crosslinked peptide pairs. Recent advances have enabled high-coverage, proteome-wide experiments that recover tens of thousands of crosslinked peptide pairs from a single sample, directly identifying thousands of pairwise protein–protein interactions with amino acid-level resolution^9–11^. XL-MS is thus a particularly valuable tool for studying protein interactions in divergent organisms that lack extensive functional annotation of their proteomes—the vast majority of species.

Here, we developed a high-coverage XL-MS strategy to map the proteome-wide interactome of *Toxoplasma gondii*, a widespread human pathogen that infects nearly a quarter of the world’s population^12,13^. *Toxoplasma* is a tractable model for the eukaryotic phylum Apicomplexa, obligate intracellular parasites separated from classical model organisms by over one billion years of evolutionary history^14,15^. The use of XL-MS in apicomplexans, including *Toxoplasma*, has been limited to the study of select protein complexes or relatively small numbers of highly abundant proteins^16–20^. Here, we identified 29,624 crosslinked peptide pairs from which we constructed a network of 2,859 pairwise protein–protein interactions, and integrated these data with predictive structural modeling to generate a proteome-scale spatial map of the parasite cell. This resource enabled the systematic annotation of protein complexes across the proteome and identified highly divergent, cryptic components of conserved eukaryotic assemblies, including RNA polymerase I and the nuclear pore complex. Most strikingly, we identified and characterized a unique ATP synthase-like complex containing a highly divergent, apicomplexan-specific α subunit that is essential for parasite fitness. Together, these findings establish proteome-wide XL-MS as a powerful and broadly applicable strategy for uncovering unexpected biology in divergent organisms, and provide a foundational resource for discovery in apicomplexan biology.

## RESULTS

### A proteome-wide XL-MS strategy for *Toxoplasma*

We designed a proteome-wide XL-MS strategy for *Toxoplasma* using the amine-reactive crosslinker disuccinimidyl sulfoxide (DSSO), whose MS-cleavable bonds facilitate identification of crosslinked peptides in complex samples^21^ **(Figure 1A)**. Intact *Toxoplasma* cells showed dramatically reduced crosslinking compared to detergent-solubilized lysate, even at high DSSO concentrations **(Figure 1B, Supplementary Figure 1A– D)**. Therefore, we used nitrogen cavitation^22^ or formaldehyde fixation followed by detergent permeabilization^23^ as sample preparation methods that enabled efficient crosslinking while preserving proteins in a near-native context.

**Figure 1.**
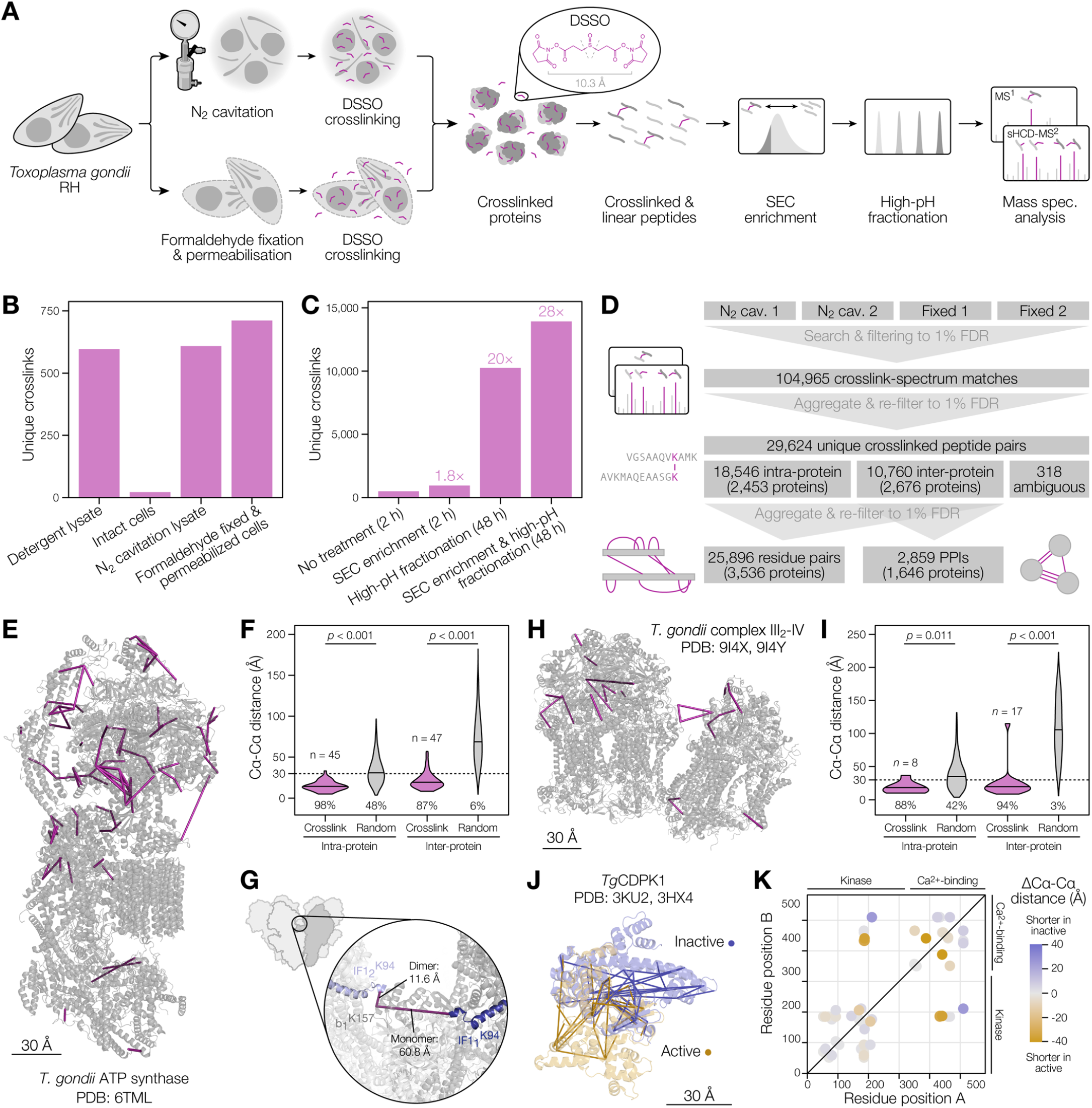
Implementation of proteome-wide XL-MS in *Toxoplasma*. **(A)** *Toxoplasma* XL-MS workflow. *T. gondii* RH tachyzoites were prepared by either nitrogen cavitation or formaldehyde fixation and permeabilization, then proximal lysine residues were crosslinked using 1 mM DSSO. Proteins were digested, and crosslinked peptides enriched by size-exclusion chromatography, followed by high-pH reversed-phase fractionation and analysis by LC–MS with a stepped-HCD MS^2^ strategy. **(B)** Number of crosslinks identified from a single 2-h LC–MS analysis of different *Toxoplasma* sample types crosslinked with 1 mM DSSO. **(C)** Effect of enrichment and fractionation strategies on the number of identified crosslinks. The total LC–MS acquisition time is noted for each strategy. (**D)** Numbers of crosslink–spectrum matches, unique crosslinks, unique residue pairs, and unique protein–protein interactions (PPIs) identified across four combined replicates. Data were refiltered at each level to maintain a 1% target–decoy false-discovery rate (FDR). **(E)** Crosslinks (magenta) mapped onto the cryo-EM structure of the *Toxoplasma* mitochondrial ATP synthase^26^. **(F)** Distribution of ATP synthase Cα–Cα distances derived from XL-MS compared to all possible lysine pairs (“random”). The proportion of Cα–Cα distances under 30 Å was tested by one-sided exact binomial test against the proportion observed among all possible intra/inter-protein lysine–lysine pairs. **(G)** Identification of a subunit b–IF1 crosslink supporting multimeric assembly of the ATP synthase. Distance of crosslinked residues is illustrated within or between monomers for comparison. **(H)** Crosslinks mapped onto the cryo-EM structure of the *Toxoplasma* mitochondrial respiratory supercomplex III_2_–IV^28^. **(I)** Distribution of supercomplex III_2_–IV Cα–Cα distances derived from XL-MS compared to all possible lysine pairs (“random”). The proportion of Cα– Cα distances under 30 Å was tested by a one-sided exact binomial test against the proportion observed among all possible intra/inter-protein lysine–lysine pairs. **(J)** Crosslinks mapped onto crystal structures of inactive and Ca^2+^-bound active CDPK1. **(K)** Changes in Cα–Cα distances of CDPK1 crosslinks between inactive and active conformations.

To increase the number of crosslink identifications, we implemented a sequential enrichment and fractionation workflow. First, crosslinks were enriched by size-exclusion chromatography based on their larger size relative to linear peptides^24^ **(Supplementary Figure 1E– G)**. Then, to increase overall proteome coverage, the enriched material was fractionated by high-resolution, high-pH, reversed-phase chromatography and pooled into 24 fractions, each analyzed by a 2 h LC–MS run using a stepped-HCD acquisition strategy^25^ **(Supplementary Figure 1H–I)**. This strategy routinely yielded >10,000 unique crosslinked peptide pairs (unique crosslinks) per sample **(Figure 1C, Supplementary Figure 1J)**.

We performed four experimental replicates: two with nitrogen cavitation and two with fixation and permeabilization. Crosslinked peptides from the fixed samples were less biased toward cytosolic proteins **(Supplementary Figure 1K)**, consistent with organelle membranes presenting a residual barrier to crosslinking after cavitation. Fixed samples also yielded slightly higher proportions of inter-protein and intra-compartment crosslinks **(Supplementary Figure 1L–M)**, consistent with better preservation of native architecture and of fragile complexes. Aggregating the data from all four replicates yielded a final dataset of 29,624 unique crosslinks at a combined crosslink-level false-discovery rate of 1% **(Figure 1D, Supplementary Data 1)**. To our knowledge, this is the largest single-crosslinker dataset generated for any organism to date.

### Crosslinks capture native protein structure and dynamics

To assess whether the identified crosslinks accurately report protein conformations, we mapped a subset onto the experimentally solved structure of the *Toxoplasma* mitochondrial ATP synthase^26^ **(Figure 1E)**. 45 intra-protein and 47 inter-protein crosslinks could be measured on this structure, of which 98% and 87% respectively fell within the expected 30 Å Cα–Cα distance limit for DSSO^27^ **(Figure 1F)**. By contrast, randomized lysine–lysine pairs in this structure only satisfied this distance constraint in 48% of intra-protein and 6% of inter-protein pairs—significantly less than the identified crosslinks in both cases (one-sided exact binomial test *p* < 0.001 for both intra- and inter-protein pairs). Notably, several crosslinks could only be rationalized by considering the higher-order hexameric assembly of ATP synthase^26^ **(Figure 1G)**. For example, we identified a crosslink between the *b* and IF1 subunits that spans >60 Å in a single ATP synthase monomer but represents only 11.6 Å when connecting two adjacent monomers. Similar results were obtained for the *Toxoplasma* mitochondrial III2–IV respiratory supercomplex^28^, with 23 of 25 crosslinked Cα–Cα pairs within 30 Å **(Figure 1H–I)**. These results confirm that our XL-MS strategy accurately captures the architecture of such protein complexes.

We next investigated whether our XL-MS data could resolve dynamic protein conformations. We mapped 30 crosslinks onto calcium-dependent protein kinase 1 (CDPK1), which undergoes a substantial structural rearrangement upon calcium binding^29^ **(Figure 1J)**. Individually, the apo and calcium-bound structures respectively satisfied 71% and 82% of crosslinks at the 30 Å Cα–Cα distance limit. However, when both structures were considered together, 87% of crosslinks could be accounted for within this limit, with three of the four remaining crosslinks marginally exceeding the limit at 33.8–35.8 Å. Comparison of crosslink distances between the two states revealed that the largest shifts mapped to the calcium-binding region **(Figure 1K)**, directly highlighting the expected conformational transition and showing that the crosslinks can capture multiple protein states coexisting in the cell^30^.

Together, these results validate the XL-MS dataset by demonstrating strong concordance between the identified crosslinks and experimentally solved protein structures, while also highlighting how crosslinks not explained by a single structure may reflect higher-order assemblies or alternative conformational states.

### XL-MS data cross-validate protein and complex structure predictions

To further evaluate the XL-MS data, we predicted protein structures using Chai-1^31^ for the 1,054 proteins within its token limit, of which 62% were predicted with high global confidence (predicted template modeling score, pTM > 0.5) **(Supplementary Figure 2A)**. 82% of intra-protein crosslinks on residue pairs with confident relative placement (mean predicted aligned error, pAE < 15) fell within the 30 Å DSSO distance limit **(Figure 2A, Supplementary Figure 2B, Supplementary Data 2A)**. By contrast, only 50% of randomized lysine–lysine pairs fell within the 30 Å window, a significantly lower proportion (one-sided Fisher’s exact test, *p* < 0.001) **(Figure 2A, Supplementary Data 2B)**, demonstrating that experimental crosslinks consistently capture residues in closer predicted proximity than random expectation. For residues modeled with low confidence (mean pAE > 15), the satisfaction rate was reduced but still exceeded that of randomized pairs (48% versus 24%, *p* < 0.001). Crosslinks in high-confidence regions were consistently satisfied across all five Chai-1 models, whereas those in low-confidence regions met the threshold in some models but not others, reflecting inconsistent residue placement **(Figure 2B)**. Together, these results demonstrate broad agreement between XL-MS data and pAE scores in assessing the local quality of predicted structures.

**Figure 2.**
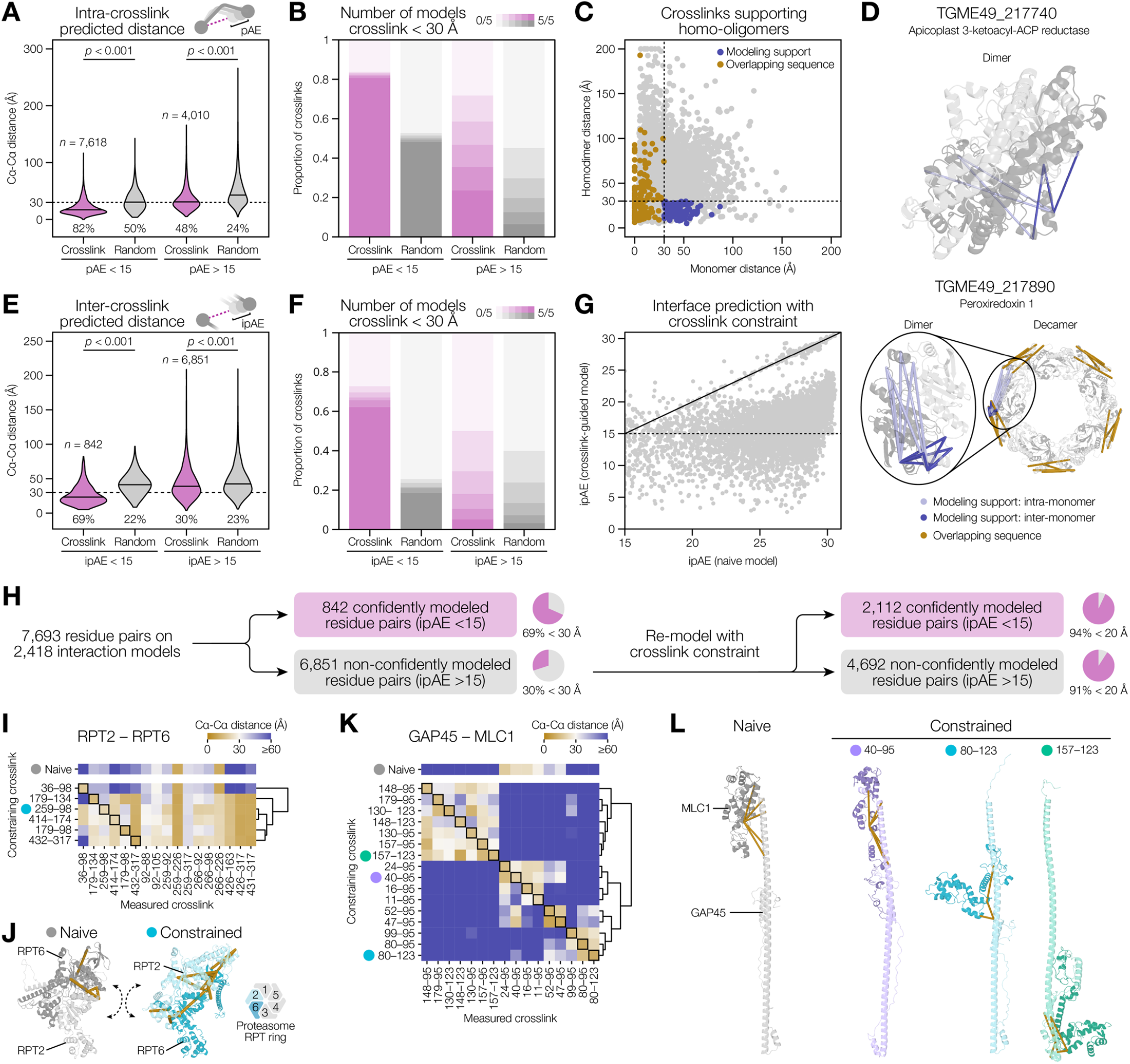
Cross-validation of XL-MS data with predicted protein structures. **(A)** Distribution of intra-protein crosslink Cα–Cα distances versus randomized lysine–lysine pairs on structures predicted using Chai-1 without crosslink constraints (naive), grouped by mean predicted aligned error (pAE). Mean pAE scores were calculated across the crosslinked residues and ±3 flanking residues. Distances are taken from the model with the lowest pAE for each residue pair. The proportion of Cα–Cα distances under 30 Å was tested by one-sided Fisher’s exact test comparing experimental crosslinks to randomized pairs within each confidence group. **(B)** Proportion of intra-protein crosslinks with Cα–Cα distances under 30 Å in 0–5 Chai-1 models, compared to randomized lysine–lysine pairs. **(C)** Comparison of intra-protein crosslink distances in naively predicted monomeric structures versus naively predicted homodimers. Crosslinks exceeding 30 Å in the monomer prediction but measured below 30 Å in the homodimer prediction with pAE < 15 are considered to support homo-oligomerization. Crosslinks with overlapping peptide sequences supporting homo-oligomerization are additionally labeled. Axes are truncated at 200 Å. **(D)** Predicted homodimer structures of TGME49_217740, an apicoplast-localized 3-ketoacyl(acyl-carrier protein) reductase homolog, and TGME49_217890, peroxiredoxin 1. The latter is additionally modeled as a homodecamer by aligning five homodimer predictions to the human peroxiredoxin 1 crystal structure (PDB: 7LJ1)^33^. Crosslinks supporting homo-oligomerization through modeling are shown as light blue (intra-molecular) and dark blue (inter-molecular) links. For peroxiredoxin-1, crosslinks with overlapping sequences are shown in gold as the shortest possible link in the decameric structure. **(E)** Distribution of inter-protein crosslink distances versus randomized lysine–lysine pairs on naive Chai-1–predicted interfaces, grouped by mean interface predicted aligned error (ipAE) of the crosslinked residues. Distances are taken from the model with the lowest ipAE for each residue pair. The proportion of Cα–Cα distances under 30 Å was tested by one-sided Fisher’s exact test comparing experimental crosslinks to randomized pairs within each confidence group.**(F)** Proportion of inter-protein crosslinks measured under 30 Å in 0–5 Chai-1 interface models, compared to randomized lysine–lysine pairs. **(G)** Comparison of ipAE for inter-protein crosslinked residues in naive (unconstrained) predictions versus predictions using the crosslink as a distance constraint at a maximum of 20 Å. **(H)** Workflow illustrating modeling of protein interfaces using crosslink constraints. **(I)** Heatmap showing crosslink distances measured in unconstrained and constrained models of proteasome subunits RPT2 and RPT6. Labels indicate crosslinked residue positions, with the first and second numbers corresponding to positions on RPT2 and RPT6, respectively. Crosslinks with low confidence (ipAE > 15) in the unconstrained model were individually applied as distance constraints; these are indicated by black boxes in the heatmap. Colored circles denote models shown in panel J. **(J)** Unconstrained and constrained models of RPT2 and RPT6. In the unconstrained model, subunits are predicted in the incorrect relative position compared to the known proteasome structure. Addition of crosslink constraints places the subunits in the correct order. Satisfied crosslinks are shown for each model. **(K)** Crosslink distances in unconstrained and constrained models of glideosome proteins GAP45 and MLC1. Labels indicate crosslinked residue positions, with the first and second numbers corresponding to positions on GAP45 and MLC1, respectively. Crosslinks with low confidence (ipAE > 15) in the unconstrained model were individually applied as distance constraints; these are indicated by black boxes in the heatmap. Colored circles denote models shown in panel L. **(L)** Unconstrained and three crosslink-constrained models of GAP45 and MLC1, revealing three binding interfaces, each satisfying a subset of crosslinks shown for each model.

Because many proteins form homo-oligomers^32^, we asked whether overlength intra-protein crosslinks in monomeric models might instead reflect higher-order assemblies. Modeling all proteins as homodimers, we identified 116 crosslinks that exceeded 30 Å as intra-monomer links but were satisfied within this limit as inter-molecular links with high confidence (mean interface predicted aligned error, ipAE < 15), supporting homo-oligomerization of 43 proteins **(Figure 2C, Supplementary Data 2C)**. These included homologs of known multimers such as isocitrate dehydrogenase, nuclear transport factor 2, and an apicoplast-localized 3-ketoacyl(acyl-carrier protein) reductase; for the latter example, three crosslinks supported homodimerization but were not satisfied in the monomer **(Figure 2D)**. As an alternative means of identifying homo-oligomers, we identified 307 intra-protein crosslinks with overlapping peptide sequences that must derive from two copies of the same protein **(Figure 2C)**. Because the crosslinked residues are close in primary sequence, they rarely exceed 30 Å distances in the monomeric structures, despite necessarily originating from homo-oligomers. Such overlapping peptides supported 142 proteins as homo-oligomers. Some proteins were supported by both approaches; a peroxiredoxin 1 homolog had 8 crosslinks that fell below 30 Å only in the homodimer model and 4 sequence-overlapping crosslinks. Together, the observed crosslinks provide strong support for higher-order assembly of the *Toxoplasma* peroxiredoxin into the decameric ring documented for the human homolog^33^ **(Figure 2D)**.

We next evaluated inter-protein crosslinks by similarly predicting structures of all interacting protein pairs using Chai-1. As with intra-protein crosslinks, the majority (69%) of inter-protein crosslinks in low-ipAE regions fell within 30 Å **(Figure 2E)** and did so consistently across the five models **(Figure 2F)**. However, comparatively few structures were modeled with high global confidence (interface predicted template modeling score, ipTM > 0.5), and only 11% of inter-protein crosslinks fell in regions of low ipAE **(Supplementary Figure 2C–D)**, consistent with other interactome-wide modeling efforts^11^.

To improve interface modeling, we re-predicted structures using individual inter-protein crosslinks that were initially modeled with low confidence as distance constraints **(Figure 2G)**. The ipAE of most crosslinked residue pairs decreased when that crosslink was included, allowing us to confidently model 2,082 additional residue pairs at ipAE < 15 **(Figure 2H)**. To assess whether constraints improved the interface model more broadly, we inspected models of interacting proteins supported by multiple inter-protein crosslinks. In some cases, a single crosslink constraint was sufficient to guide prediction of a model that satisfied all other crosslinks where the unconstrained model had failed. An example is the interaction between RPT2 (PSMC1, TGME49_267080) and RPT6 (PSMC5, TGME49_261010), adjacent subunits of the heterohexameric ATPase ring of the proteasome 19S regulatory particle. These proteins are homologous, and the naive (unconstrained) prediction placed them in the wrong relative orientation, leaving most crosslinks unsatisfied. Five of six crosslinks used as restraints guided the subunits into the correct orientation, yielding a model consistent with biochemical studies and solved structures in other eukaryotes^34,35^ **(Figure 2I–J)**. By contrast, for other protein pairs, no single model satisfied all crosslinks simultaneously, even when constraints were applied. For the glideosome proteins GAP45 (TGME49_223940) and MLC1 (TGME49_257680)^36^, the pattern of crosslink satisfaction across constrained models suggested multiple binding modes, each accounting for a distinct subset of identified crosslinks **(Figure 2K–L)**.

Together, these analyses demonstrate how XL-MS data and artificial intelligence-guided structural predictions can cross-validate each other, while also revealing the limitations of current prediction models. For both monomers and heterodimers, identified crosslinks on confidently predicted residues are largely satisfied within the expected distance limit. Yet although the majority of monomeric structures can be predicted confidently, this is rarely true of heterodimeric interfaces. We show that incorporation of crosslink constraints can guide predictions toward correct models but also uncover complexities arising from alternative interaction interfaces.

### The XL-MS protein interaction network captures cellular organization from complexes to organelles

We investigated the organization of the protein interaction network constructed from the inter-protein crosslinks, comprising 2,859 protein–protein interactions among 1,646 proteins at a 1% interaction-level FDR **(Figure 3A, Supplementary Data 1)**. Of note, this network encompassed 69% (118/171) of interactions reported at the same FDR in a prior XL-MS study of *Toxoplasma*^19^, while expanding the total interactome size nearly 17-fold. To evaluate the global network structure, we examined the distribution of interactions **(Figure 3B)**. Protein–protein interaction networks are expected to follow a power-law distribution, in which most proteins have few connections and a small subset acts as highly connected hubs^37^. The distribution of our network was best fit by a power-law model (α = 1.73), which achieved the lowest Akaike Information Criterion among alternative models (geometric, Poisson, Weibull, log-normal, and negative binomial) and passed a Kolmogorov–Smirnov goodness-of-fit test (bootstrap *p* = 0.84)^38^. These findings support a topology consistent with established properties of biological interaction networks. Highly connected proteins exhibited lower ratios of non-synonymous to synonymous nucleotide substitutions (dN/dS) **(Figure 3C)**, consistent with purifying selection acting on central hubs to preserve network stability^39^. However, the distribution of interactions may also reflect technical limitations of XL-MS, as crosslinks are more readily detected for more abundant proteins, which tend toward lower dN/dS ratios **(Figure 3D)**.

**Figure 3.**
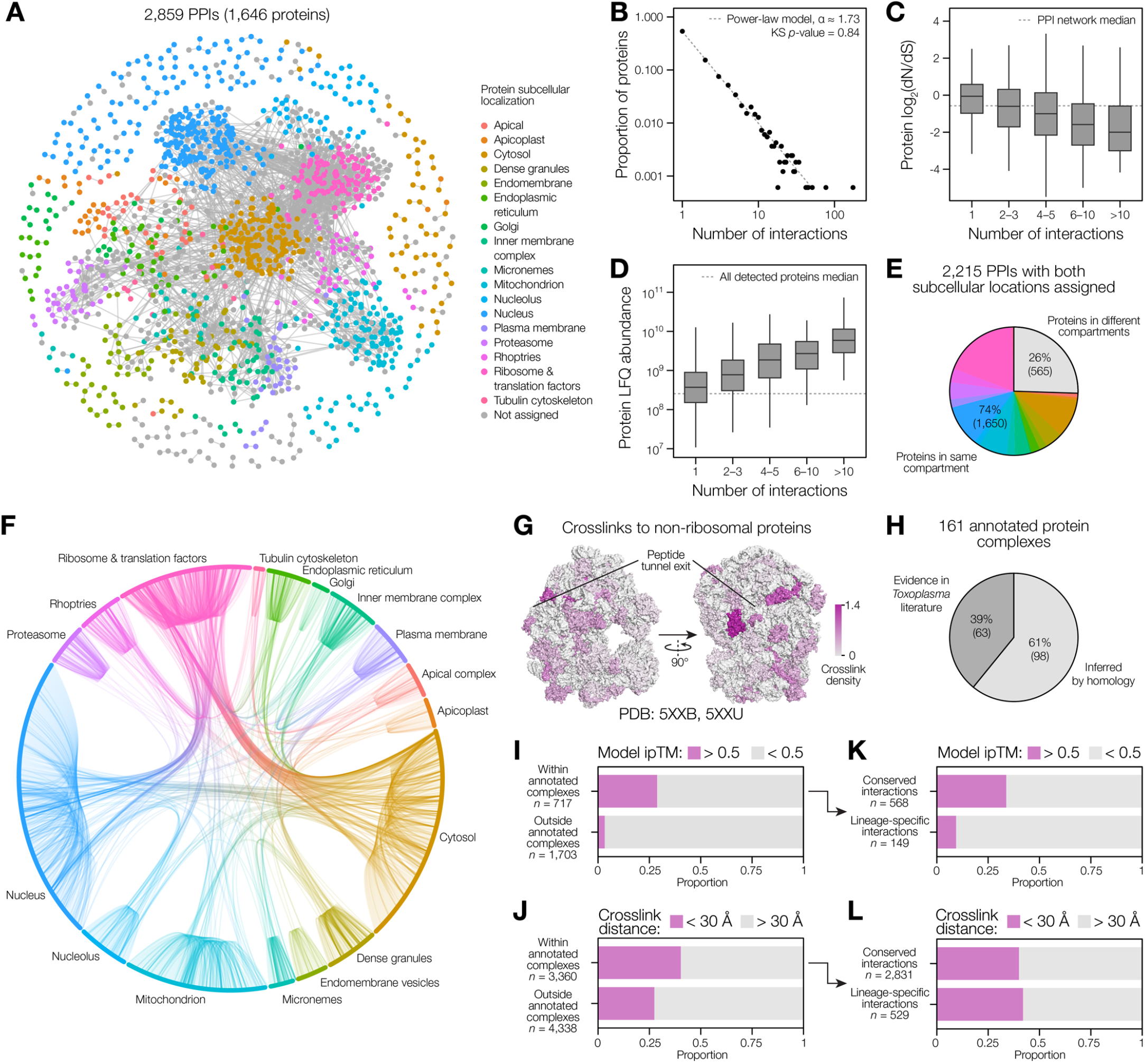
The XL-MS–derived protein interaction network recapitulates cellular organization. **(A)** Global protein–protein interaction network derived from inter-protein crosslinks, filtered to a 1% interaction-level FDR. **(B)** Distribution of PPI network node degrees (number of interactions per protein). The dotted line illustrates a power-law distribution with an estimated α (scaling exponent) of 1.73. The degree distribution of the PPI network did not differ significantly from the power-law model (Kolmogorov–Smirnov test, *p* = 0.84). **(C)** Ratio of non-synonymous to synonymous single-nucleotide polymorphisms (dN/dS) for proteins in the PPI network, grouped by number of interactions. dN/dS data were obtained from ToxoDB^88^. The dotted line represents the median of all proteins in the network. **(D)** Label-free quantification (LFQ) abundance of proteins in the network, grouped by number of interactions. Abundance values were determined from the XL-MS experiments by taking the geometric mean across all four experiments. The dotted line represents the median abundance of all detected proteins, regardless of whether crosslinks were detected for them. **(E)** Proportion of interactions for proteins with assigned subcellular locations that occur within the same or different subcellular compartment. Same-compartment interactions are colored by compartment using the same color scheme as in panel A. **(F)** Schematic illustration of interactions between proteins with assigned subcellular locations. **(G)** Density of crosslinks between the ribosome and other compartments mapped onto the structure of the *Toxoplasma* ribosome^89^. Ribosomal proteins are colored by the number of inter-compartment crosslinks divided by the total number of lysine residues in the protein. **(H)** Proportion of annotated protein complexes within the protein interaction network that have direct experimental evidence in *Toxoplasma* or are inferred by homology to characterized complexes in other organisms. **(I)** Proportion of inter-protein crosslinks modeled under 30 Å in predicted interface structures, grouped by whether the proteins are in the same annotated complex. **(J)** Proportion of protein interfaces modeled with an ipTM score > 0.5, grouped by whether the proteins are in the same annotated complex. **(K)** Proportion of protein interfaces within annotated complexes modeled with an ipTM score > 0.5, grouped by protein conservation. Interactions between proteins that are conserved eukaryote-wide, as defined by OrthoMCL ortholog groups^88^, are classified as conserved, whereas those involving one or more lineage-specific proteins are classified as lineage-specific. **(L)** Proportion of crosslinks within annotated complexes modeled at a distance under 30 Å in predicted interface structures, grouped by protein conservation, defined as above.

To assess whether the network reflects native cellular organization, we assigned subcellular localizations to the proteins in the network using prior spatial proteomics data^40^ supplemented with manual curation **(Supplementary Data 3)**. Of the interactions for which both proteins had assigned localizations, 74% linked proteins within the same compartment **(Figure 3E)**. By contrast, randomizing interactions while maintaining the same number of contacts per protein yielded a network where only 14% of interactions occurred within the same compartment. These results demonstrate that the XL-MS network faithfully recapitulates the spatial architecture of the cell and performs far above random expectation.

The majority of inter-compartment interactions involved the ribosome and its associated translation factors **(Figure 3F)**. We hypothesized that many of these interactions reflect transient proximity during translation, as nascent chains of any compartment will briefly encounter ribosomal proteins. Consistent with this interpretation, inter-compartment crosslinks were strongly enriched around the peptide exit tunnel, with the three most crosslinked proteins clustering in this region and the most crosslinked single residue mapping to lysine 98 on the solvent-exposed face of uL29 adjacent to the tunnel **(Figure 3G)**. However, several biologically meaningful inter-compartment interactions were also apparent. Some interactions reflected ribosomal biogenesis and function, connecting to nucleolar processome complexes and the endoplasmic reticulum (ER) Sec61 translocon. Other inter-compartment interactions linked physically or functionally related structures: the apical complex and tubulin cytoskeleton, ER and Golgi, and inner membrane complex and plasma membrane. A final subset reflected dually localized proteins, including nucleocytoplasmic transport receptors^41^ and a VCP/p97 homolog that interacted with nuclear spindle disassembly^42^ and ER-associated degradation components^43^.

Having established that the XL-MS data largely recapitulate subcellular organization, we next examined whether they could resolve the assembly of individual protein complexes. We systematically annotated protein complexes across the global PPI network, identifying 161 complexes, of which 63 were supported by direct experimental evidence in *Toxoplasma* and 98 were inferred by homology to characterized complexes in other organisms **(Figure 3H, Supplementary Data 3)**. Together, these complexes accounted for 37% (1,055/2,859) of all detected PPIs and 48% (5,142/10,760) of all inter-protein crosslinks. Protein pairs within the same complex were more likely to be modeled with high confidence (ipTM > 0.5) **(Figure 3I)**, and their crosslinks were more frequently satisfied at the 30 Å distance limit for DSSO **(Figure 3J)**. Higher model confidence was generally correlated with protein conservation, reflecting the greater availability of solved homologous structures in the prediction training data **(Figure 3K)**. By contrast, crosslink distance satisfaction within annotated complexes did not differ between conserved and lineage-specific proteins, indicating that crosslinking validates interactions independently of the evolutionary conservation that biases structural prediction **(Figure 3L)**.

These analyses establish that XL-MS data can capture subcellular organization, supporting over 160 known protein complexes—many previously identified in *Toxoplasma* and others inferred solely by homology— and broadly validating the network’s ability to recover known biology. Additionally, our curation revealed numerous uncharacterized proteins associated with well-defined complexes, positioning them as candidate novel subunits.

### Targeted validation of interaction networks identifies cryptic components of essential eukaryotic complexes

To evaluate whether the network accurately identified novel protein interactions, we selected uncharacterized proteins associated with conserved eukaryotic protein complexes for targeted experimental validation. We focused on the combined nuclear and nucleolar interaction networks as the most extensive group of complexes in our dataset **(Figure 4A, Supplementary Figure 3)**.

**Figure 4.**
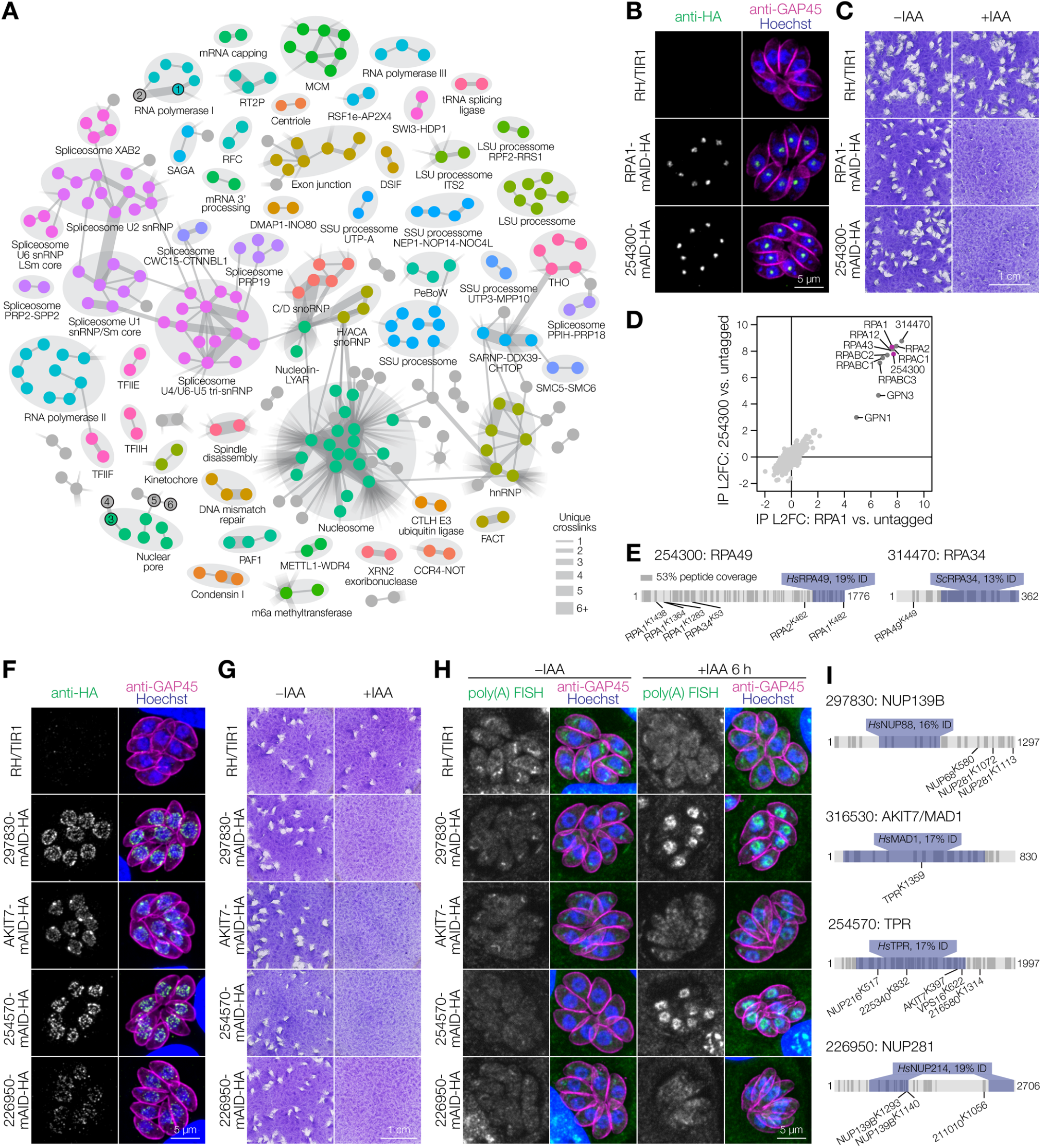
XL-MS identifies novel components of conserved protein complexes. **(A)** XL-MS-derived nuclear and nucleolar protein interactome. Proteins are represented as nodes and interactions as connections between them. The width of each connection is proportional to the number of unique crosslinks supporting each interaction. Proteins further investigated in this figure are labeled: 1 = RPA1, 2 = 254300, 3 = NUP139B, 4 = 226950, 5 = 254570, 6 = AKIT7. Refer to **Supplementary Figure 3** for full annotation. **(B)** Immunofluorescence microscopy of endogenously tagged RNA polymerase I subunit RPA1 and candidate subunit 254300. **(C)** Plaque assays for RPA1 and 254300 inducible knockdown cell lines treated with vehicle (–IAA) or auxin (+IAA) to induce knockdown. The parental RH/TIR1 cell line is included as a control. **(D)** Anti-HA co-immunoprecipitation–mass spectrometry of RPA1 and 254300. Data show log_2_-transformed fold-change of DIA protein abundance relative to an untagged control (*n* = 3 replicates per strain). All labeled proteins were significantly enriched for both RPA1 and 254300 compared to control (L2FC ≥ 2; *p*_adj_ ≤ 0.01, one-sided *t*-test with Benjamini–Hochberg false-discovery rate correction). **(E)** Schematics of 254300 and 314470 indicating regions of homology identified using HHpred (blue), peptide coverage from co-immunoprecipitation experiments (shaded regions), and the positions of identified crosslinks. **(F)** Immunofluorescence microscopy of endogenously tagged candidate nuclear pore complex (NPC) proteins. **(G)** Plaque assays for candidate NPC protein inducible knockdown cell lines treated with vehicle (–IAA) or auxin (+IAA) to induce knockdown. The parental RH/TIR1 cell line is included as a control. **(H)** RNA fluorescence in situ hybridization with a poly(dT) probe and immunofluorescence microscopy of candidate NPC protein inducible knockdown cell lines left untreated (–IAA) or following 6 h of knockdown (+IAA). **(I)** Schematics of candidate nuclear pore complex proteins indicating regions of homology identified using HHpred (blue), peptide coverage from combined XL-MS experiments (shaded regions), and the positions of identified crosslinks.

First, we identified an uncharacterized protein with strong evidence for interaction with RNA polymerase I subunits RPA1 and RPA2: TGME49_254300 (hereafter referred to by its numeric identifier). As the large size of both proteins prevented structural modeling, this interaction represented an unbiased test case for experimental validation. We endogenously tagged 254300 and RPA1 with an HA epitope and mini auxin-inducible degron (mAID) for rapid, inducible protein depletion^44^ **(Supplementary Figure 4A–D)**. By immunofluorescence microscopy, we determined that both proteins localized to a nuclear punctum with reduced DNA staining consistent with the nucleolus—the expected location of RNA polymerase I **(Figure 4B)**. Plaque assays, which measure overall *Toxoplasma* growth, showed that depletion of either RPA1 or 254300 disrupted parasite replication, in line with the expected essential role of RNA polymerase I and prior genome-wide CRISPR knockout data^45^ **(Figure 4C)**. Co-immunoprecipitation–mass spectrometry demonstrated reciprocal enrichment of RPA1 and 254300, validating their interaction **(Figure 4D, Supplementary Data 4)**. In addition, both proteins co-enriched an identical set of interactors comprising all other annotated RNA polymerase I subunits, GPN1, GPN3, and an additional uncharacterized protein, TGME49_314470.

Although GPN1 and GPN3 have not been studied in *Toxoplasma*, they mediate nuclear import of RNA polymerase II in humans and may therefore have wider functions encompassing RNA polymerase I in other organisms^46^. Analysis of 254300 with HHpred^47^ identified a C-terminal region with remote homology to RPA49, preceded by an apicomplexan-specific N-terminal extension of approximately 1,500 amino acids **(Figure 4E)**. The gene model was supported by 53% peptide coverage across the length of the protein. Notably, most crosslinks between 254300 and other RNA polymerase I components mapped to this apicomplexan-specific N-terminal region, indicating its involvement in the architecture of the complex. Similarly, HHpred identified the additional uncharacterized protein 314470 as a putative homolog of RPA34 **(Figure 4E)**. The stringent re-filtering needed to maintain a 1% interaction-level FDR had excluded a single identified crosslink between 314470 and 254300 from the final interaction network; however, RPA49 and RPA34 form a heterodimeric subcomplex within RNA polymerase I^48^, and given the homology-based identities assigned above, the excluded crosslink is consistent with an equivalent arrangement in *Toxoplasma*.

We next investigated the nuclear pore complex (NPC), which is structurally divergent in *Toxoplasma* and remains poorly characterized^49^. We endogenously tagged four putative members of this complex: TGME49_297830 (identified as *Tg*NUP139B during the course of this work^50^), TGME49_316530 (a homolog of *Plasmodium falciparum* AKIT7^51^), TGME49_254570, and TGME49_226950 **(Supplementary Figure 4E–I)**. All four proteins localized to the nuclear periphery, consistent with the expected location of the NPC **(Figure 4F)**. AKIT7 additionally accumulated at a single perinuclear focus, consistent with its expected kinetochore localization^51^. All four proteins were shown to be essential by plaque assay following mAID-mediated depletion **(Figure 4G)**.

As these proteins could not be solubilized under non-denaturing conditions for co-immunoprecipitation, we instead sought functional evidence for their involvement in NPC activity by assessing export of polyadenylated mRNA using RNA fluorescence in situ hybridization **(Figure 4H)**. Following a 6-h knockdown of 297830 (TgNUP139B) or 254570, we observed accumulation of poly(A) mRNAs within the parasite nucleus, but did not observe clear accumulation following depletion of AKIT7 or 226950.

Finally, we assessed these proteins for remote homology to conserved eukaryotic NPC components **(Figure 4I)**. 297830 (TgNUP139B) was identified as a homolog of human NUP88. As previously reported, AKIT7 is a homolog of MAD1^51^, a spindle assembly checkpoint protein that associates with the NPC in humans and yeast^52,53^. We identified 254570 as a homolog of nucleoprotein TPR, whereas 226950 has partial homology to *Hs*NUP214. As *Hs*NUP214 and its yeast homolog *Sc*NUP159 are required for export of mRNAs from the nucleus^54,55^, the lack of phenotype for 226950 in *Toxoplasma* mRNA export suggests functional divergence over evolutionary time.

Together, these analyses demonstrate that XL-MS– derived interaction networks accurately recapitulate known protein complexes and can be used to reliably identify uncharacterized components of essential macromolecular complexes, establishing XL-MS as a powerful framework for hypothesis generation and system-scale analysis of the *Toxoplasma* interactome.

### A divergent apicomplexan α subunit defines a novel ATP synthase subcomplex

In the mitochondrial interaction network, we identified an uncharacterized protein associated with the ATP synthase β subunit that is absent from the solved ATP synthase structure and was not detected in previous co-immunoprecipitation or complexome profiling studies^26,56–58^ **(Figure 5A, Supplementary Figure 5)**. This interaction was modeled with moderate confidence (Chai-1 ipTM = 0.66), yet the crosslink identified between the two proteins exceeded the distance limit for DSSO suggesting the need for further structural refinement **(Supplementary Data 2)**. Sequence analysis identified this uncharacterized protein as a highly divergent homolog of the ATP synthase α subunit. This α-like protein, which we designate α2, is present only in *Toxoplasma* and related coccidians, as well as in the photosynthetic relatives of apicomplexans, *Chromera velia* and *Vitrella brassicaformis*, suggesting it was present in the common ancestor of apicomplexans and chromerids but lost in many apicomplexan lineages **(Figure 5B)**. Although α2 shares only 25% sequence identity with the canonical *Toxoplasma* α subunit (hereafter α1)—compared with 71% between α1 and human α—the two paralogs are predicted to have highly similar core folds **(Figure 5C)**. In contrast to α1, which is under strong purifying selection, α2 shows neutral to modestly diversifying selection, consistent with functional divergence between the two paralogs **(Figure 5D)**.

**Figure 5.**
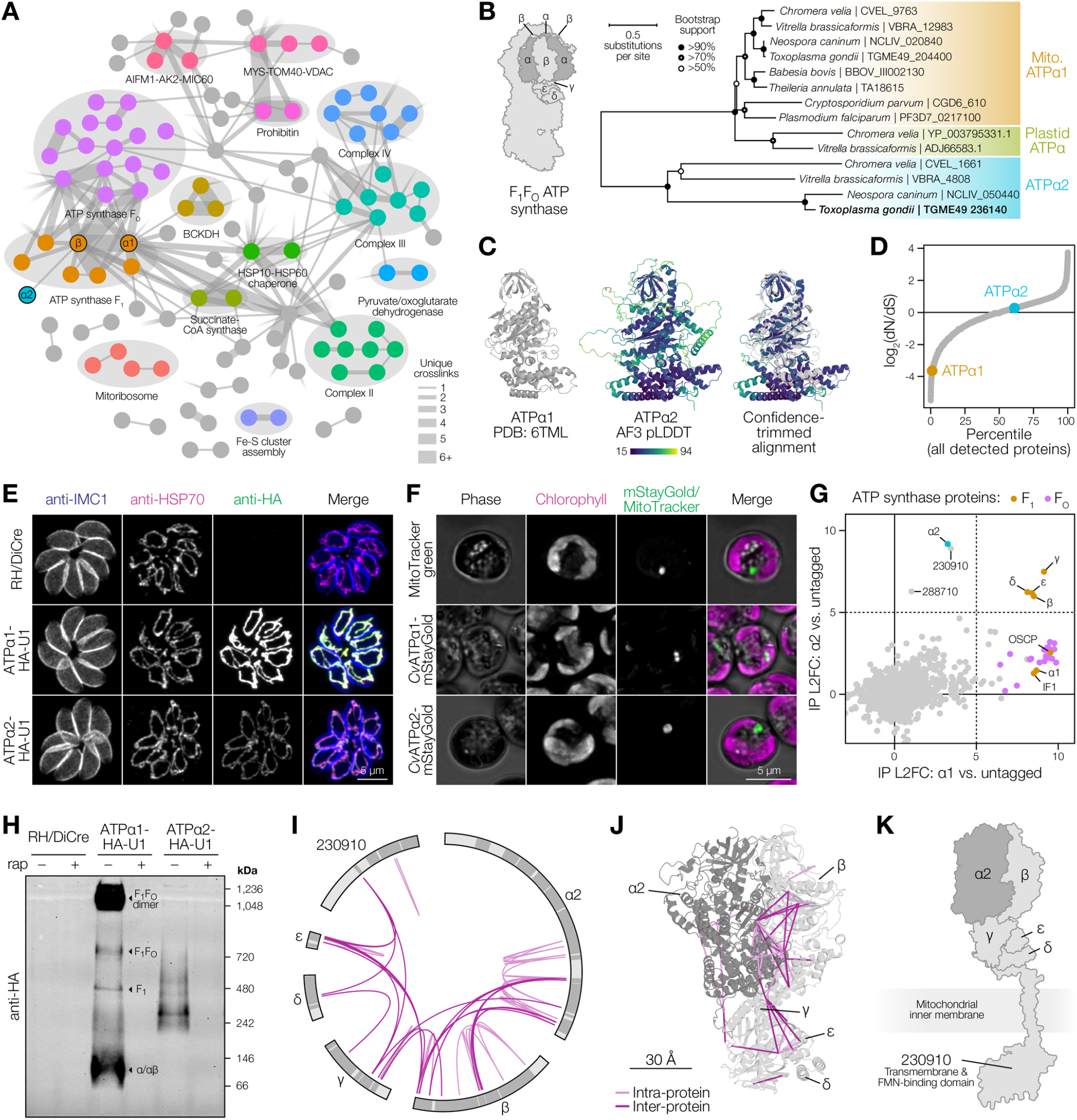
Identification of a variant mitochondrial ATP synthase subcomplex. **(A)** XL-MS-derived mitochondrial protein interactome. Proteins are represented as nodes and interactions as connections between them. The width of each connection is proportional to the number of unique crosslinks supporting each interaction. Proteins further investigated in this figure are labeled. Refer to **Supplementary Figure 4** for full annotation. **(B)** Maximum-likelihood phylogeny of select apicomplexan and chromerid ATP synthase α subunits. **(C)** Cryo-EM structure of *Toxoplasma* ATPα^26^, AlphaFold 3 prediction of ATPα2 colored by pLDDT, and structural alignment with low-confidence surface loops removed. **(D)** Percentile-ranked non-synonymous to synonymous SNP ratios across all *Toxoplasma* isolates^88^ for all proteins detected in the XL-MS dataset. **(E)** Immunofluorescence microscopy of endogenously tagged *Toxoplasma* α homologs. **(F)** Live fluorescence microscopy of *Chromera velia* stained with MitoTracker green or transiently expressing mStayGold-tagged α homologs. **(G)** Anti-HA co-immunoprecipitation–mass spectrometry of ATPα1 and ATPα2 in *Toxoplasma*. Data show log_2_-transformed fold-change of DIA protein abundance relative to an untagged control (*n* = 6 for untagged and ATPα2, *n* = 3 for ATPα1). All detected ATP synthase subunits are highlighted. **(H)** Immunoblot of native PAGE probed with anti-HA for endogenously tagged *Toxoplasma* ATPα1 and ATPα2 with and without rapamycin-induced, U1-mediated knockdown. **(I)** Crosslinks detected among ATPα2 complex members following targeted crosslinking–immunoprecipitation–mass spectrometry. Protein sequences are illustrated from N-to C-terminus in clockwise orientation. Shaded regions of proteins indicate peptide coverage across all XL-MS and co-immunoprecipitation experiments. Light pink connections indicate intra-protein crosslinks while dark pink connections indicate inter-protein crosslinks. **(J)** Chai-1 prediction of the ATPα2 complex structure using all detected inter-protein crosslinks as modeling constraints with a maximum distance limit of 35 Å. **(K)** Schematic representation of the predicted ATPα2 complex structure including the uncharacterized protein 230910.

To verify whether α2 is a bona fide ATP synthase interactor and further explore its properties, we generated conditional knockdown strains for both α paralogs. We endogenously tagged both α1 and α2 with an HA epitope and a synthetic loxP-flanked 3′ UTR followed by U1 binding sites, in a DiCre-expressing *Toxoplasma* strain. The tagged alleles could be conditionally depleted upon rapamycin treatment^59^ **(Supplementary Figure 6A–D)**. Immunofluorescence microscopy confirmed that both α1 and α2 localize to the *Toxoplasma* mitochondrion; however, α2 was expressed at approximately one-twentieth the level of α1 **(Figure 5E)**. Using a recently developed transfection system for *Chromera velia*^60^, we observed that mStayGold-tagged homologs of α1 and α2 localized to spherical structures distinct from the plastid and similar in appearance to compartments labeled by MitoTracker **(Figure 5F)**. Limited transfection efficiency precluded colocalization of the tagged α homologs with MitoTracker labeling in *Chromera*. However, heterologous expression of the *Chromera* α homologs in *Toxoplasma* showed that both were targeted to the mitochondrion, supporting conserved localization across the two evolutionarily distinct species **(Supplementary Figure 6E)**.

To compare α1 and α2 interacting partners, we performed co-immunoprecipitation–mass spectrometry using the tagged subunits **(Figure 5G, Supplementary Figure 6F, Supplementary Data 5)**. Assessing enrichment at a log2 fold-change cutoff of 5, both α1 and α2 co-purified with the β, γ, δ, and ε subunits of the ATP synthase F1 subcomplex, whereas OSCP, IF1, and the FO subunits were enriched only with α1. α2 additionally co-purified with two uncharacterized mitochondrial proteins (TGME49_230910 and TGME49_288710). Notably, α1 and α2 did not strongly co-enrich with each other, indicating that they assemble predominantly into distinct complexes.

Consistent with its distinct subunit composition, the α2-containing complex migrated at ∼250–300 kDa by native PAGE, distinct from both the complete F1FO ATP synthase (monomer ∼720 kDa; dimer ∼1 MDa) and the detached F1 (∼480 kDa) observed for canonical α **(Figure 5H)**. As α2 is approximately double the molecular weight of α1 **(Supplementary Figure 6D)**, yet the α2-containing complex migrates below the canonical F1, this result indicates a stoichiometry different from the canonical α3β3γδε arrangement.

To further define the architecture of the α2 complex, we performed targeted crosslinking– immunoprecipitation using DSSO, identifying 61 unique crosslinks **(Figure 5I, Supplementary Figure 6G, Supplementary Data 6)**. Of the 25 detected inter-protein links, 12 were between α2 and β and one between α2 and γ, supporting direct contact of these proteins. The complete set of inter-protein constraints was used to guide structure prediction with Chai-1^31^ assuming a single copy of each subunit (α2, β, γ, δ, ε), approximating the molecular weight estimated for the complex by native PAGE. The resulting model suggested an overall organization resembling the canonical F1 core, with all α2–β crosslinks supporting a single interface geometry similar to the catalytic ADP-binding interface of F1, but incorporating two additional α2-specific helices **(Figure 5J)**. The position of one extended helix would sterically preclude IF1 binding, providing a structural rationale for the absence of IF1 from the α2 complex.

Crosslinks also supported interactions with the uncharacterized mitochondrial protein TGME49_230910, which is predicted to contain an unstructured N terminus followed by a transmembrane helix and a flavin mononucleotide–binding domain. Crosslinks mapped the unstructured region of TGME49_230910 to the γ, δ, and ε subunits, suggesting a potential anchoring role analogous to the FO domain **(Figure 5K)**. No crosslinks were detected between TGME49_288710 and other proteins, precluding positioning of this protein within the complex.

Together, these data identify a distinct mitochondrial ATP synthase subcomplex containing the highly divergent, apicomplexan-specific α2 subunit. This complex forms an F1-like assembly with altered subunit stoichiometry, associates with at least one additional uncharacterized protein, but does not associate with the FO domain, suggesting divergence from the canonical function of the mitochondrial ATP synthase.

### The α2 complex regulates ATP synthase stability and parasite fitness

To determine whether α2 is required for *Toxoplasma* fitness, we examined the effect of knockdown in the endogenously tagged strains^59^ **(Figure 6A)**. Plaque assays following knockdown of α2 revealed severely reduced parasite growth **(Figure 6B)**, though parasites remained viable in culture for at least two weeks after depletion. By contrast, no parasite growth was observed following knockdown of α1, indicating that, while both paralogs are required for full parasite fitness, they may have distinct functions.

**Figure 6.**
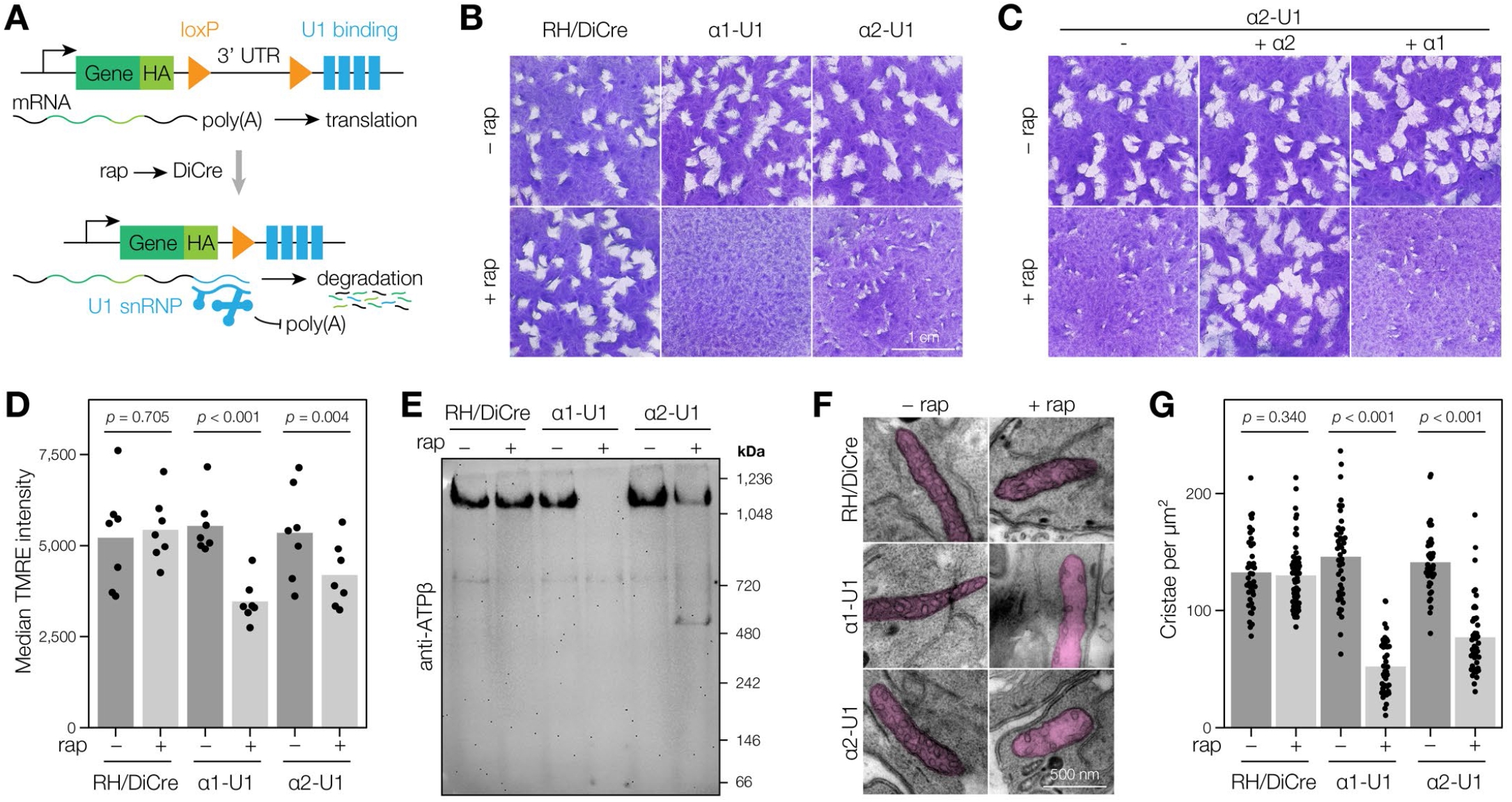
Distinct roles of α1 and α2 in mitochondrial ATP synthase regulation. **(A)** Schematic of U1-mediated inducible protein knockdown. **(B)** Plaque assays for ATPα1 and ATPα2 inducible knockdown cell lines treated with vehicle (–rap) or rapamycin (+rap) to induce knockdown. The parental RH/DiCre cell line is included as a control. **(C)** ATPα2 inducible knockdown cell line and ATPα2/ATPα1 complemented cell lines treated with vehicle (–rap) or rapamycin (+rap) to induce knockdown. **(D)** Measurement of mitochondrial membrane potential via TMRE fluorescence by flow cytometry following knockdown of ATPα1 or ATPα2. Statistical significance was assessed by one-sided paired *t*-test (*n* = 5). **(E)** Immunoblot of native PAGE probed with anti-ATPβ following rapamycin-induced, U1-mediated knockdown of ATPα1 and ATPα2. **(F)** Transmission electron micrographs of the *Toxoplasma* mitochondrion following rapamycin-induced, U1-mediated knockdown of ATPα1 and ATPα2. The mitochondrion is indicated by pink shading. For additional images, refer to **Supplementary Figure 8. (G)** Quantification of mitochondrial cristae density from transmission electron micrographs following knockdown of ATPα1 and ATPα2. Each parasite was considered as a replicate with at least 40 parasites analyzed per condition. Statistical significance was assessed by one-sided *t*-test.

To determine whether α1 and α2 are functionally distinct, we introduced a copy of either α1 or α2 into the α2 conditional knockdown strain at a neutral locus^61^, with expression driven by the α2 promoter to achieve expression levels comparable to those of the endogenous α2 copy **(Supplementary Figure 7A–E)**. Complementation with α2 rescued parasite growth in a plaque assay, whereas expression of α1 at equivalent levels did not, demonstrating that α2 performs a role that cannot be fulfilled by the canonical α subunit **(Figure 6C)**.

To investigate the mechanistic basis of this difference, we first examined mitochondrial membrane potential as a broad indicator of mitochondrial health. Measurement of TMRE fluorescence by flow cytometry^62^ revealed that knockdown of either α1 or α2 resulted in a decreased mitochondrial inner membrane potential (ΔΨm) **(Figure 6D)**. For the canonical α1 subunit, this is consistent with a prior report in *Toxoplasma* that disruption of the ATP synthase results in depolarization^63^. The similar effect observed upon α2 depletion suggests that the α2 complex is also required for ΔΨm maintenance, but does not distinguish whether this is mediated through the canonical ATP synthase or an independent mechanism.

While examining complexes by native PAGE, we noticed that knockdown of α2 reduced the amount of intact F1FO complex concomitant with the appearance of a ∼480 kDa band consistent with a dissociated F1 subcomplex, indicating that α2 function is required for the stable association of the F1 and FO modules **(Figure 6E)**. This phenotype was distinct from depletion of α1, for which we observed complete loss of high molecular weight complexes. Given that ATP synthase oligomers are known to play a structural role in shaping mitochondrial cristae^26^, we examined mitochondrial ultrastructure by transmission electron microscopy. Knockdown of α2 resulted in loss of cristae architecture, mirroring the phenotype observed upon depletion of α1 and confirming the loss of ATP synthase integrity **(Figure 6F–G, Supplementary Figure 8)**.

Together, these data demonstrate that, although the α2 complex is distinct in structure and composition from the canonical ATP synthase, it is required for stable F1FO assembly. Destabilization of the ATP synthase upon α2 depletion is accompanied by loss of cristae architecture and collapse of ΔΨm, accounting for the fitness defect observed in its absence. The inability of canonical α1 to compensate for this role shows that the α2 complex has acquired a non-redundant function in mitochondrial physiology, marking a unique adaptation of this lineage.

## DISCUSSION

We present a comprehensive protein interactome for the parasite *Toxoplasma gondii*, derived from deep-coverage crosslinking mass spectrometry. The dataset accurately captures protein structures and can be leveraged to model protein complexes. We demonstrate that it reliably identifies novel protein–protein interactions, enabling the discovery of divergent components of RNA polymerase I and the nuclear pore complex. We also define a novel mitochondrial ATP synthase-like complex, characterizing a divergent, lineage-specific α subunit that regulates ATP synthase stability, maintenance of mitochondrial membrane potential, and parasite fitness. In total, we show that this dataset is a powerful resource for discovery in apicomplexan biology.

Creation of this dataset required two technical improvements in the XL-MS workflow. First, sample preparation by nitrogen cavitation^22^ or fixation and permeabilization^23^ overcame the poor membrane permeability of DSSO, providing a general strategy for crosslinking of recalcitrant samples. Second, extensive high-resolution fractionation by high-performance liquid chromatography dramatically increased crosslink coverage, as these species are inherently low in abundance and disproportionately benefit from deeper sampling. Together with size exclusion chromatography^24^ and careful validation of each step in the workflow, these advances yielded a robust pipeline that reproducibly identified over 10,000 crosslinks per sample. Across experimental replicates, we identified over 29,000 crosslinks, supporting a network of over 2,800 protein– protein interactions—the largest XL-MS dataset for a single crosslinker to date. Importantly, this network was obtained after stringent filtering to a 1% interaction-level FDR, which is critical for confidence in the data given the hierarchical structure of XL-MS data^64^ but is often not applied in comparable datasets. Constructing a network of this scale would require a campaign of thousands of individual co-immunoprecipitation or proximity labeling experiments, an undertaking feasible only for the most heavily resourced consortia^1,2,65^. Yet XL-MS offers greater precision than other proteome-wide approaches such as hyperLOPIT^40^, co-fractionation profiling^56,66^, or computational prediction^67,68^, as the detected analyte is a physical artifact that maps proximal sites on each partner.

The spatial resolution of XL-MS data can be leveraged in modeling protein structures and complexes. Artificial intelligence has transformed structure prediction, yet training data for protein–protein interactions remain biased toward stable complexes from select model organisms, and large-scale predictions lack empirical validation at comparable scales^68,69^. Our crosslinks agree remarkably well with empirically determined and confidently predicted structures, yet a large gap remains in modeling protein interfaces compared to monomeric proteins—even for experimentally validated interacting pairs—underscoring the limitations of current predictions. Beyond validation, XL-MS data provide orthogonal, empirical constraints from near-native contexts to guide structure prediction, and are already being incorporated into modeling frameworks^31,70,71^. Consistent with previous work^70^, we find that crosslink-derived distance constraints can improve model confidence, yet also highlight complexities that future frameworks will need to accommodate. In some cases, crosslinks drive the model toward a single satisfying structure, while in others, they capture multiple protein states, homo-oligomeric assemblies, or alternative binding modes^30^. Here, we deliberately used single crosslink restraints so as not to coerce incorrect structures; future efforts may benefit from careful incorporation of multiple restraints. Crucially, refining structural models at this resolution brings us closer to identifying functionally relevant interfaces that can be disrupted chemically or investigated experimentally in more targeted ways than the deletion of entire subunits.

A major application of this dataset is the identification of novel protein complex components, including de-orphaning of hypothetical proteins lacking prior functional annotation. The scale of discovery is evident in the capture of at least 161 protein complexes, the majority previously unstudied in *Toxoplasma*. We illustrate the accuracy of these inferences with targeted validation of two essential complexes. Our interaction network identified two uncharacterized proteins, TGME49_254300 and TGME49_314470, as components of RNA polymerase I, which we confirmed by co-immunoprecipitation and subsequently assigned as orthologs of RPA49 and RPA34, respectively. The RPA34/RPA49 heterodimer is the least conserved component of RNA polymerase I across eukaryotes^48^, with exceptional sequence divergence likely explaining why these proteins were previously thought absent in Apicomplexa^72^. A similar theme was apparent for the nuclear pore complex, for which we identified three divergent components with remote homology to conserved NPC-associated proteins that had evaded detection by co-immunoprecipitation or proximity labeling^50,73,74^. In situ cryo-electron microscopy has shown that the *Toxoplasma* NPC has an unusual architecture, apparently lacking the cytoplasmic Y-complex and nuclear basket^49^. Our identification of a divergent TPR homolog—the primary scaffold of the basket—suggests this structure does exist but may be flexible or dynamically associated with the pore, as in yeast^75^, and thus not resolved by cryo-electron microscopy. Sequence and ultrastructural divergence appears to be coupled to functional divergence: we identified a conserved subcomplex comprising *Tg*NUP281 (TGME49_226950) and *Tg*NUP139B (TGME49_297830), which are homologs of *Hs*NUP214/*Sc*NUP159 and *Hs*NUP88/*Sc*NUP82, respectively^76^. While *Tg*NUP139B is required for mRNA export, *Tg*NUP281 appears dispensable, in contrast to the yeast and human homologs^54,55^. These two complexes represent only a fraction of the novel biology accessible in this dataset, underscoring the power of XL-MS to systematically identify components of poorly characterized complexes in divergent organisms.

The interaction data can also reveal entirely novel complexes, as exemplified by the α2 complex we characterize here. The α and β subunits of F-type ATP synthases are among the most conserved proteins across bacteria and eukaryotes, yet organisms rarely maintain functionally distinct paralogs of these subunits. In *Toxoplasma*, however, we identified a divergent α paralog (α2) that participates in a mitochondrial complex distinct from the canonical ATP synthase. By combining proteome-wide and targeted XL-MS, co-immunoprecipitation–mass spectrometry, and crosslink-informed structural modeling, we propose an architecture for this complex that resembles the core F1 domain but differs in composition and stoichiometry. Although depletion of either α subunit produces similar phenotypes—disruption of mitochondrial ultrastructure and membrane potential—α2 is not functionally interchangeable with the canonical α1. This functional distinction raises two possibilities. First, the α2 complex may play a role in ATP synthase assembly or regulation, processes that differ substantially across organisms and compartments and remain poorly characterized in apicomplexans^77,78^. Alternatively, it may serve an independent function whose loss produces similarly broad mitochondrial defects. To our knowledge, no comparable complex has been described in any other organism, highlighting how even core bioenergetic machinery can harbor undiscovered lineage-specific innovations.

*Toxoplasma* is a highly tractable model for the Apicomplexa, enabling the functional investigations presented here, yet the integrated proteome-wide XL-MS and structural modeling framework we establish is applicable to any culturable organism, including those that are not genetically tractable. The study of diverse organisms illuminates how core cellular processes have evolved under distinct selective pressures; this approach provides a uniquely powerful structural perspective through which to investigate these questions across the tree of life. Moreover, emerging AI-driven tools for antimicrobial drug design increasingly depend on accurate structural models, which remain largely unavailable for non-model pathogens; proteome-wide XL-MS offers a scalable means of generating the experimental constraints needed to fill this gap. More broadly, this framework may be extended to complex host–pathogen interaction systems, where molecular interface data are of direct therapeutic relevance. Finally, adapting XL-MS for quantitative, proteome-wide comparisons^79^ will open an entirely new perspective on cellular organization, enabling systematic mapping of how protein interaction networks are remodeled across cellular states.

## MATERIALS & METHODS

### Cell culture

#### Mammalian cell lines

Human foreskin fibroblasts (HFFs) and A7r5 cells were cultured in DMEM (Gibco #11965118) supplemented with 10% heat-inactivated fetal bovine serum (Sigma #F4135) and 2 mM L-glutamine. Cells were grown in a humidified incubator at 37°C and 5% CO_2_.

##### Toxoplasma gondii

*Toxoplasma gondii* strains as detailed in the Key Resources Table **(Supplementary Data 7)** were maintained by serial passage in HFFs in DMEM supplemented with 3% heat-inactivated newborn calf serum (Sigma #N4762), 2 mM L-glutamine, and 10 µg/mL gentamicin. Protein knockdown via the mAID system was induced by addition of 50 µM indole-3-acetic acid (IAA), which was maintained for the duration of the experiment. Knockdown via the U1 system was induced by treatment with 50 nM rapamycin for 2 h, followed by washout. Unless otherwise stated, U1-mediated knockdown was induced 96 h prior to the start of experimental assays.

##### Chromera velia

*Chromera velia* strain CCMP2878 was cultured in L1 medium without SiO_3_ (Bigelow Laboratory for Ocean Sciences) in an environmental chamber (Percival Scientific) at 27°C with a 14 h light/10 h dark cycle.

### Proteome-wide crosslinking mass spectrometry

#### Sample preparation and crosslinking

10–15 × 15 cm dishes of freshly egressed *T. gondii* RHΔ*KU80* tachyzoites were used per experiment. The parasites were mechanically released from remaining host cells, passed through a 5 µm filter to remove host cell debris, and washed twice with PBS to remove residual protein and amino acids from the culture medium.

For crosslinking optimization using detergent lysate, the cells were lysed on ice for 10 min in 2 mL lysis buffer (1% IGEPAL CA-630, 150 mM NaCl, 50 mM HEPES·NaOH pH 7.4, 1× cOmplete Mini EDTA-free protease inhibitor) and clarified by centrifugation at 21,000 × *g*. Lysate protein concentration was determined by BCA assay and adjusted with additional cavitation buffer to obtain a 2 mg/mL final concentration during the crosslinking reaction. Freshly prepared 20 mM DSSO in anhydrous DMSO was added to a final concentration of 1 mM, unless otherwise specified. The crosslinking reaction was incubated at 25°C for 30 min with constant agitation and quenched by the addition of 1 M ammonium bicarbonate to a final concentration of 50 mM for a further 15 min. Following the crosslinking reaction, SDS was added to 5% v/v to denature the crosslinked proteins.

For nitrogen cavitation, the cells were resuspended in 2 mL chilled cavitation buffer (250 mM sucrose, 50 mM HEPES·NaOH pH 7.4, 1× cOmplete Mini EDTA-free protease inhibitor). The cell suspension was pressurized with nitrogen to 2000 psi in a chilled Parr Instruments 4639 Cell Disruption Vessel and allowed to equilibrate on ice for 20 min. The system was depressurized via the release valve, causing cell disruption as nitrogen bubbles form within the cells. The lysate was collected via the release valve and centrifuged to remove remaining intact cells. Lysate protein concentration was determined by Bradford assay and adjusted with additional cavitation buffer to obtain a 2 mg/mL final concentration during the crosslinking reaction. DSSO crosslinking at 1 mM and quenching proceeded as described above, followed by the addition of SDS to 5% v/v.

For crosslinking of fixed cells, the cells were resuspended in 20 mL 4% w/v methanol-free formaldehyde in PBS to fix for 15 min at room temperature with occasional agitation. 20 mL 0.2% Triton X-100 in PBS was added for a final concentration of 0.1% for 10 min to permeabilize the cells. The cells were washed twice with PBS and resuspended in 2.4 mL PBS. DSSO crosslinking at 1 mM and quenching proceeded as described above. Following crosslinking, the cells were pelleted and resuspended in 5% v/v SDS, 50 mM HEPES·NaOH pH 7.4 and heated at 95°C for 2 h to reverse the formaldehyde fixation.

#### Protein digestion

Crosslinked proteins were digested using a modified SP3 protocol^80^ as follows. The sample was diluted to approximately 1 mg protein/mL with 50 mM triethylammonium bicarbonate (TEAB). Tris(2-carboxyethyl)phosphine and 2-chloroacetamide were added to final concentrations of 10 mM and 40 mM respectively and the sample was heated at 70°C for 30 min or 95°C for 10 min to reduce disulfide bonds and alkylate cysteine residues. 0.1× sample volume of an equal mix of Sera-Mag Carboxylate-Modified E3 and E7 Magnetic Beads was added, followed by 4× volumes of ethanol or acetonitrile to precipitate proteins onto the beads. The sample was rotated at room temperature for 10 min, after which the beads were bound to a magnet and washed three times with 80% ethanol. The beads were resuspended in 1 mL 50 mM TEAB with trypsin/Lys-C protease mix at a 1:100 protease:sample mass ratio. The protein sample was digested at 37°C for 16 h with constant agitation. The resulting peptides were desalted using a Pierce Peptide Desalting Spin Column, quantified by Pierce Quantitative Fluorometric Peptide Assay, and lyophilized.

#### Size-exclusion chromatography

The crosslinked peptide sample was resuspended in SEC mobile phase (30% v/v acetonitrile, 0.1% v/v trifluoroacetic acid in water) to approximately 10 mg/mL. 25 µL of peptide sample was injected onto a Superdex 30 Increase 3.2 × 300 mm column (Cytiva) attached to an ÄKTA pure 25 chromatography system (Cytiva) and separated by isocratic elution at a flow rate of 50 µL/min over two column volumes. Two-minute fractions were collected. The concentration of peptides in each fraction was quantified by Pierce Quantitative Fluorometric Peptide Assay. The earliest three fractions containing an above-baseline concentration of peptides, representing the top ∼10% of the sample by size and exceeding 1.5-fold enrichment of CSMs as a proportion of total spectrum matches, were pooled and lyophilized. SEC fractionation was repeated with additional material to accumulate at least 50 µg of crosslink-enriched peptides.

#### High-pH reversed-phase fractionation

The crosslink-enriched peptides were fractionated using a 1260 Infinity II high-performance liquid chromatography system (Agilent) with an XBridge Peptide BEH C18 3.5 µm particle size, 130 Å pore size, 2.1 mm × 250 mm column (Waters). The sample was taken up at approximately 1 µg/µL and 50 µL were injected onto the column. Separation was conducted with a mobile phase consisting of 0.0175% v/v ammonium hydroxide, 0.01125% v/v formic acid, 2% v/v acetonitrile in water (solution A) and 0.0175% v/v ammonium hydroxide, 0.01125% v/v formic acid, 80% acetonitrile in water (solution B) at a flow rate of 300 µL/min. The column temperature was held constant at 25°C and separation was monitored with a multiple wavelength detector at 205, 254, and 280 nm. The column was conditioned with 10 column volumes of 80% solution B followed by 10 column volumes of 0% B. The sample was separated over a linear gradient of 0–6% B over 9 min, 6–28.5% B over 61 min, 28.5–34% B over 3.5 min, 34–40% B over 13.5 min, and a wash of 40% B for 8 min, for a total length of 96 min. One-minute fractions were collected resulting in 96 fractions of 300 µL, which were concatenated every 24th fraction to produce 24 superfractions that were lyophilized and resuspended in 0.2% v/v formic acid prior to LC–MS analysis.

#### Liquid chromatography–mass spectrometry

LC–MS analysis was performed with a Vanquish Neo nanoLC system coupled to an Orbitrap Eclipse Tribrid mass spectrometer equipped with an EASY-Spray electrospray ionization source and FAIMS Pro interface (all Thermo Fisher Scientific). NanoLC separation used an Acclaim PepMap C18 75 µm × 2 cm trap column (Thermo Fisher Scientific) combined with a Generation 4 Aurora series Ultimate UHPLC C18 1.7 µm particle size, 250 mm × 75 µm analytical column (IonOptiks) with a mobile phase consisting of 0.1% v/v formic acid in water (solution A) and 0.1% v/v formic acid, 80% v/v acetonitrile in water (solution B) at a flow rate of 400 nL/min and column temperature held constant at 50°C. 500 ng of each superfraction was separated over a linear gradient of 3–40% solution B over 100 min, 40–95% solution B over 10 min, and a wash of 95% solution B for 6 min.

The mass spectrometer was operated in positive mode with the ion source temperature set to 305°C. Spectra were acquired with a 2 s cycle time, alternating between FAIMS Pro compensation voltages of −50/−60/−75 V for each cycle. MS^1^ Orbitrap spectra were acquired at a resolution of 120,000, scan range of 375–1500 m/z, standard automatic gain control settings, and automatic ion injection time. To select precursors for fragmentation, the monoisotopic peak selection filter was used in peptide mode, the intensity filter was used with a threshold of 2.5 × 10^4^, the charge state filter was used to select precursors with a charge of 3–8, and the dynamic exclusion filter was used with an exclusion duration of 30 s. MS^2^ Orbitrap spectra were acquired in DDA mode with an isolation window of 1.6 m/z and a resolution of 30,000 using a stepped-HCD fragmentation with normalized collision energies of 21%, 27%, and 33%. Standard AGC target was used with a maximum ion injection time of 100 ms.

#### Crosslink, peptide, and protein identification

Crosslinked peptides were identified from mass spectra using XlinkX^81,82^ via Proteome Discoverer 2.4 (Thermo Fisher Scientific) using the “MS^2^” acquisition strategy for DSSO. MS^2^ spectra were first partitioned on the presence of the paired fragment peaks diagnostic of DSSO cleavage: spectra containing these peaks at a signal-to-noise ratio of 1.5 or greater were directed to the crosslink search with XlinkX, and all remaining spectra to the linear peptide search with Sequest HT. Putative crosslink spectra were searched against the *Toxoplasma gondii* RH88 strain proteome (retrieved from ToxoDB release 68), allowing crosslinking of lysine residues and protein N-termini with up to two missed cleavages. Cysteine carbamidomethylation was set as a static modification and methionine oxidation was set as variable modification. Crosslink–spectrum matches (CSMs) with an identification score greater than 20 and Δscore greater than 10 were accepted, which we empirically determined to result in a target-decoy FDR of ∼1%. Non-crosslink MS^2^ spectra were searched against the same reference proteome using Sequest HT additionally allowing hydrolyzed DSSO (+176.014 Da) and ammonium-quenched DSSO (+175.030 Da) as variable modifications on lysine residues. Target-decoy FDR filtering with a strict cutoff of 1% and relaxed cutoff of 5% was applied at all levels to peptide-spectrum matches, unique peptides, and proteins.

#### Crosslink data processing

Target and decoy CSMs were exported from the Proteome Discoverer “Crosslink MS^2^ Scans” and further processed using custom R scripts. CSMs were aggregated to unique crosslinked peptide pairs, retaining the highest CSM identification score and corresponding Δscore as the crosslink score. Following aggregation, identification score and Δscore cutoffs were adjusted to maintain a 1% crosslink-level target-decoy FDR while maximizing the number of target crosslinks. Crosslinked peptides mapping to more than one protein were disambiguated using parsimony-based principles analogous to protein inference in standard proteomics as follows: (1) protein accessions with no uniquely identifying crosslinked peptides were discarded from consideration; (2) since intra-protein crosslinks are more probable than inter-protein crosslinks, crosslinks comprising one ambiguous and one non-ambiguous peptide were assigned as an intra-protein crosslink of the non-ambiguous accession where this was a possibility; (3) manually curated protein homolog groups were collapsed by assigning crosslinked peptides to the most abundant group member (e.g., GAPM1A, TGRH88_036510, was selected over GAPM1B, TGRH88_036500). Eliminated protein accessions for each crosslinked peptide are noted in the “Unique Crosslinks” supplementary table. The remaining 318 crosslinks containing ambiguous peptides that could not be resolved in the above steps were discarded from downstream analyses. The peptide crosslinks were aggregated to unique residue-residue pairs, retaining the highest crosslink identification score and corresponding Δscore as the residue pair score. Following aggregation, identification score and Δscore cutoffs were again adjusted to maintain a 1% residue pair-level target-decoy FDR. In parallel, the peptide crosslinks were aggregated to protein–protein interactions by summing the identification scores and Δscores of all contributing crosslinks.

Informed by the resulting distribution of target and decoy identification scores and Δscores, we applied an additional minimum identification score or minimum Δscore filter to achieve a 1% PPI-level target-decoy FDR.

### Chai-1 modeling

Structure predictions were performed for all heterodimers identified by XL-MS, and each unique interacting protein was predicted as both a monomer and homodimer. We used Chai-1 because it was, at the time of this study, the only available method with an AlphaFold 3-style architecture supporting implicit residue-level crosslink modeling, and for its speed and single-sequence performance.

The predictions were made using a local installation of the publicly available pretrained Chai-1 model by Chai Discovery^31^ on an NVIDIA GH200 Grace Hopper Superchip (96 GB GPU HBM, 525 GiB CPU system memory); this code was modified to save and export the predicted aligned error (pAE) matrices without altering the model inference. All predictions were performed using the default settings of 3 recycles, 200 diffusion time steps, and 5 diffusion output samples with a fixed random seed for reproducibility. Due to the limited and uneven homolog depth for *Toxoplasma*, multiple sequence alignments (MSAs) were not used at inference; however, predictions were made using ESM embeddings. Predictions of naive heterodimers, monomers, and homodimers were made without any defined restraints. The total XL-MS dataset coverage was limited to predictions within the 2,048 Chai-1 token limit.

To evaluate PPI structural improvement with crosslinks, single heterodimer re-predictions were made for each unique inter-protein crosslink. These restraints were provided at inference via Chai-1’s “contact” restraint format, in which the crosslinked residue index and number were defined for each interacting protein, and contact restraint confidence was defined at 1.0. During Chai-1 training, distance thresholds were randomly sampled from 0–30 Å, leaving restraints at ≥ 30 Å under-represented. In a subset of PPI predictions, we observed that restraints defined above 20 Å are not meaningfully enforced at inference; the maximum distance was therefore set to 20 Å for all predictions, to ensure the crosslink restraint was applied.

For all five models of each unique monomer prediction, metrics were recorded for Cα Euclidean distance between all unique monomer associated intra-protein crosslink residue pairs, pAE and pTM scores, and crosslink-specific pAE scores. Similarly, for all five model outputs for PPI predictions, metrics were calculated for the Cα Euclidean distance between all inter-protein crosslink residue pairs, monomer pAE scores, overall ipAE and ipTM scores, and a crosslink-specific ipAE score. To reduce the effect of the crosslink restraint itself artificially lowering the pAE and ipAE scores at the restrained residues, we calculated these crosslink-specific scores as the mean value across the crosslinked residue pair and a ± 3-residue flanking window, averaged between the symmetric regions of the pAE matrix. For each unique PPI and crosslink pair, the model with the lowest ipAE score was used for plotting.

Null measurements were computed between random lysine pairs for the naive monomer and heterodimer predictions. Lysines were randomly sampled with replacement across all available intra-protein lysines and inter-protein lysines on monomers and multimers, respectively. To ensure comparable sampling, each PPI was assigned a number of random lysine pairs equal to the number of experimentally recorded, PPI-specific crosslinks.

Following naive heterodimer prediction, the unique PPI crosslink measures on the naive predictions were filtered for multimeric confidence using the crosslink-specific ipAE score and within expected range crosslink distances, in which ipAE < 15 Å was defined as a confident multimer and Cα distance ≤ 30 Å was defined as “within range” crosslink measurement. All unconfident PPI crosslinks, regardless of crosslink distance, were individually used to re-predict the PPI with a single crosslink restraint. These re-predicted single crosslink heterodimers were assessed using the same metrics as the naive heterodimers.

### *Toxoplasma* genetic manipulations

#### Inducible knockdown strains

Inducible knockdown strains were established using a high-throughput tagging strategy^83^. For each target, a sequence block was designed containing a sgRNA targeting the 3′ UTR of the gene and homology arms for in-frame C-terminal integration. This block was assembled into a vector containing the tagging payload via Gibson assembly. For mini auxin-inducible degron (mAID)-mediated knockdown^44^, the payload comprised mAID and an HA epitope, followed by the 3′ UTR of CDPK1 and a DHFR-TS* expression cassette for pyrimethamine selection. Approximately 10 µg of the payload vector was linearized with BsaI and transfected into RH/ΔKU80/TIR1 parasites along with 10 µg of a Cas9 expression plasmid by electroporation as previously described^83^. For U1-mediated knockdown^59^, the payload contained an HA epitope and loxP-flanked 3′ UTR of CDPK1, followed by four U1 snRNP binding sites and a DHFR-TS* expression cassette. This vector was similarly linearized with BsaI and transfected into RH/ΔKU80/DiCre parasites^84^ along with a Cas9 expression plasmid by electroporation. Following transfection, parasites were selected for integration of the payload plasmid with 1 µM pyrimethamine. Clonal lines were obtained by serial dilution and verified by diagnostic PCR.

#### Complementation strains

The promoter and 5′ UTR of ATPα2 (TGRH88_041450/TGME49_236140) and the coding sequences of ATPα2 and ATPα1 (TGRH88_034710/TGME49_204400) were amplified from RH/DiCre genomic DNA by PCR. These sequences were assembled into a vector containing a C-terminal V5 epitope tag, an HXGPRT expression cassette for mycophenolic acid/xanthine selection, an sgRNA expression cassette with a protospacer targeting neutral locus 1^61^, and homology arms for integration at neutral locus 1. A linear repair template was amplified from the plasmid by PCR and transfected into RH/DiCre/ATPα2-HA-U1 parasites together with 10 µg of a Cas9 expression plasmid by electroporation. Beginning 24 h post-transfection, parasites were selected with 25 µg/mL mycophenolic acid and 50 µg/mL xanthine. Clonal lines were obtained by serial dilution and verified by diagnostic PCR.

### *Chromera* plasmid design

Synthesis and assembly of DNA fragments were carried out by GenScript (Piscataway, NJ, USA). A *Pst*I 5′-flanked and *Bgl*II 3′-flanked DNA fragment containing the Kozak sequence, CDS of the target gene, and a linker was synthesized and ligated into the *Pst*I and *BglI*I sites of the pCvGAPDH-CvH2B-mStayGold plasmid previously described for *Chromera velia*^60^. The VEuPathDB accession numbers for the targeted genes are CVEL_9763 (*Cv*ATPα1) and CVEL_1661 (*Cv*ATPα2).

### ATPα phylogeny

ATPα homologs from apicomplexans and chromerids were identified from orthogroup OG7_0004945 using OrthoMCL. Incomplete sequences were excluded, yielding 128 taxa, and plastid ATPα sequences from *Chromera velia* (YP_003795310.1) and *Vitrella brassicaformis* (ADJ66583.1) were added. Protein sequences were aligned using MUSCLE in MEGA12 and manually trimmed to remove poorly aligned regions. Phylogenetic relationships were inferred using maximum likelihood in MEGA12^85^ under the LG+G+I model selected by the Bayesian Information Criterion. Node support was assessed using bootstrap resampling with 500 replicates.

### Fluorescence and immunofluorescence microscopy

#### *Toxoplasma* immunofluorescence microscopy—Lourido lab

HFFs cultured in 96-well high-performance glass-bottom plates (Cellvis) were infected with parasites for 24 h. Cells were fixed with 4% (w/v) formaldehyde in PBS for 15 min, permeabilized with 0.1% (v/v) Triton X-100 in PBS for 15 min, and blocked with 2% (w/v) bovine serum albumin (BSA) in PBS for 30 min. The cells were incubated with primary and secondary antibodies diluted as detailed in the Key Resources Table, for 1 h each. Staining with 0.5 µg/mL Hoechst 33258 was performed simultaneously with secondary antibody incubation. Samples were imaged on a CSU-W1 spinning-disk confocal microscope (Nikon) running NIS Elements AR using a 100×/1.35 NA plan apochromat silicone oil objective. Z-stacks were acquired with 0.2–0.3 µm spacing between optical sections. Maximum intensity projections were generated using Fiji^86^.

#### *Toxoplasma* transient transfection and immunofluorescence microscopy*—*Hu lab

RH/ΔKU80 parasites were transfected with *Chromera velia* expression plasmids as described above (*Toxoplasma* genetic manipulations) and inoculated onto A7r5 cells grown in 35 mm diameter glass-bottom dishes (MatTek). At 24 h post-transfection, the cells were fixed with 3.7% (w/v) formaldehyde in PBS for 10 min, permeabilized with 0.5% (v/v) Triton X-100 for 15 min, and blocked with 1% (w/v) bovine serum albumin for 30–60 min. The cells were incubated with primary and secondary antibodies diluted as detailed in the Key Resources Table. Images were acquired using a DeltaVision OMX Flex imaging system (GE Healthcare-Applied Precision) with a 60×/1.42 NA oil immersion lens using immersion oil with a refractive index of 1.522.

#### *Chromera* transient transfection and fluorescence microscopy— Hu lab

Freshly saturated *Chromera* cultures (OD_600_ ≈ 1.5–2.0) were diluted 1:2–1:3 in fresh medium and incubated for 4–6 days, until reaching an OD_600_ of 0.8–1.3. *Chromera* expression plasmids were linearized with *Not*I-HF and *Kpn*I-HF (NEB), ethanol precipitated, and combined with 2.5 × 10^6^ *Chromera* cells in 50 µL MAX Efficiency Transformation Reagent for Algae in a 1 mm gap electroporation cuvette. The cells were transfected using a NEPA21 electroporator (Bulldog Bio) with the following parameters: poring pulse—300 V, two pulses, 10 ms pulse length, 50 ms interval, 20% decay rate with polarity switching; transfer pulse—25 V, 10 pulses, 50 ms length, 50 ms interval, 40% decay rate with polarity switching. Immediately after electroporation, 100 µL recovery medium (0.5 M sorbitol in L1 medium without SiO_3_) was added to the cuvette and the culture was recovered with an additional 750 µL recovery medium in the dark for 24 h before returning to normal culture conditions. The cells were imaged 4–6 days post-transfection using a DeltaVision OMX Flex imaging system as described above.

#### *Chromera* MitoTracker labeling and fluorescence microscopy— Hu lab

Due to the barrier posed by the *Chromera* cell wall, MitoTracker labeling was carried out by electroporation. 2.5 × 10^6^ *Chromera* cells, prepared as described above, were resuspended in 50 µL MAX Efficiency Transformation Reagent for Algae supplemented with 0.5 µM MitoTracker Green FM and electroporated as described above. The cells were immediately resuspended in 200 µL recovery medium, incubated in the dark for 15–30 min, washed twice with recovery medium and transferred to 35 mm diameter glass-bottom dishes and imaged on a DeltaVision OMX Flex imaging system as described above.

### RNA fluorescence in situ hybridization

HFFs cultured in 96-well high-performance glass-bottom plates (Cellvis) were infected with parasites for 24 h. mAID-mediated knockdown was induced with IAA 6 h prior to fixation. Cells were fixed with 4% (v/v) formaldehyde in PBS for 15 min and permeabilized with 70% (v/v) ethanol for 10 min. 200 nM poly-d(T)_40_–Cy5 oligonucleotide was hybridized to samples overnight at 37°C using Stellaris RNA FISH buffers (Biosearch Technologies) according to the manufacturer’s protocol. Samples were subsequently immunostained and imaged on a spinning-disk confocal microscope as described above.

### Plaque assays

200–300 parasites were inoculated onto confluent HFFs in a 6-well plate and left undisturbed for 10 days. The cells were washed with PBS, fixed with ethanol for 10 min, and allowed to air dry. The cells were stained with crystal violet (50 mM crystal violet, 20% ethanol, 0.8% ammonium oxalate) for 10 min, washed 3× with PBS, and scanned on an Epson Perfection V800 Photo scanner.

### Co-immunoprecipitation–mass spectrometry

For each replicate, parasites were harvested from a single 15 cm dish of HFFs infected for 48 h, mechanically released, filtered through 5-µm membranes, and washed twice with PBS. Parasites were lysed on ice for 10 min in 1 mL IP buffer (1% IGEPAL CA-630, 150 mM NaCl, 50 mM HEPES·NaOH pH 7.4, 1× cOmplete Mini EDTA-free protease inhibitor, 0.1% benzonase) and clarified by centrifugation at 21,000 × *g*. Lysates were incubated with 50 µL anti-HA magnetic beads (Pierce) for 2 h at 4°C, washed three times with IP buffer, and eluted in 20 µL 5% SDS, 50 mM TEAB.

Eluted proteins were processed using an S-Trap digestion protocol^87^. Samples were reduced and alkylated with 10 mM TCEP and 40 mM chloroacetamide at 95°C for 10 min, followed by acidification to 2.5% (v/v) phosphoric acid. Proteins were precipitated by addition of six volumes of S-Trap buffer (100 mM TEAB–H_3_PO_4_ pH 7.55, 90% methanol), captured on S-Trap Micro spin columns (ProtiFi), and washed four times with 150 µL S-Trap buffer. On-column digestion was performed overnight at 37°C in a humidified incubator with 1 µg trypsin/Lys-C protease mix (Pierce) in 20 µL 50 mM TEAB. Peptides were sequentially eluted with 40 µL 50 mM TEAB, 40 µL 0.2% formic acid, and 40 µL 50% acetonitrile, 0.2% trifluoroacetic acid, and dried prior to LC–MS analysis.

LC–MS analysis was performed with a Vanquish Neo nanoLC system coupled to an Orbitrap Eclipse Tribrid mass spectrometer equipped with an EASY-Spray electrospray ionization source and FAIMS Pro interface, as described above. Samples were separated over a linear gradient of 3–25% solution B over 45 min, 25–35% solution B over 15 min, 35–95% B for 5 min, and a wash of a 95% solution B for 6 min. The mass spectrometer was operated in positive mode with the ion source temperature set to 305°C and FAIMS Pro compensation voltage set to −50 V. MS^1^ Orbitrap spectra were acquired with a resolution of 120,000, scan range of 350–2000 m/z, standard automatic gain control setting, and automatic ion injection time. MS^2^ Orbitrap spectra were acquired in DIA mode with a resolution of 30,000 and scan range of 375–1200 m/z, custom AGC target of 1000 units, and 30% normalized CID. Isolation windows of 25 m/z were used with 0.5 m/z overlaps.

Data were analyzed using DIA-NN v1.8.1 in library-free mode. An in silico spectral library was generated from the *Toxoplasma gondii* RH88 strain proteome (retrieved from ToxoDB release 68) by deep learning–based prediction, with tryptic digestion allowing up to two missed cleavages. Peptide charge states of 2–6 were considered, while peptide length, precursor m/z range, and fragment m/z range were left as defaults. Cysteine carbamidomethylation was set as a fixed modification and methionine oxidation as a variable modification, with N-terminal methionine excision enabled. A two-pass search strategy was employed, in which a spectral library was first created from the DIA runs and then used to reanalyze them. Results were filtered at a 1% false discovery rate. Log2 fold-changes were calculated from the geometric mean of abundance values for each protein in each sample group, and p-values were calculated by two-sided Student’s *t*-test on log-transformed abundance values with Benjamini–Hochberg false-discovery rate correction.

### Crosslinking–immunoprecipitation

RH/DiCre/ATPα2-HA-U1 parasites were harvested from 10 × 15 cm dishes of HFFs infected for 48 h and lysed in 1 mL IP buffer as described for co-immunoprecipitation–mass spectrometry. The sample was crosslinked with 1 mM DSSO for 30 min at 25°C and quenched with 50 mM ammonium bicarbonate for 15 min. ATPα2-HA was captured with 250 µL anti-HA magnetic beads, eluted, and digested by S-Trap as described above. Peptide yield was quantified by fluorometric assay (Pierce). The sample was analyzed on an Orbitrap Eclipse Tribrid mass spectrometer equipped with a FAIMS Pro source and coupled to a Vanquish HPLC system, using a stepped-HCD MS^2^ acquisition strategy as described for proteome-wide crosslinking mass spectrometry. The sample was initially analyzed using a 2 h gradient with internal FAIMS compensation voltage stepping at −50/−60/−75 V, followed by three 3 h gradients each acquired with a single compensation voltage at −50, −60, or −75 V. Data were processed using XlinkX within Proteome Discoverer, with downstream analysis performed using custom scripts as described for proteome-wide crosslinking mass spectrometry.

### SDS-PAGE and immunoblot

Parasites were mechanically released from host cells, passed through a 5 µm filter to remove host cell debris, washed twice with PBS, and lysed in buffer containing 1% (v/v) IGEPAL CA-630, 150 mM NaCl, 50 mM HEPES·NaOH (pH 7.4), and 1× Halt protease inhibitor cocktail without EDTA (Thermo Fisher) on ice for 10 min. Insoluble material was removed by centrifugation at 21,000 × *g* for 5 min at 4°C. Protein concentration was determined using the Pierce BCA Protein Assay Kit (Thermo Fisher). Proteins were separated on a 4–15% gradient Mini-PROTEAN TGX gel (Bio-Rad) at 160 V for 40 min and transferred to a nitrocellulose membrane using a Trans-Blot Turbo Transfer System (Bio-Rad). Membranes were blocked with 5% (w/v) skim milk powder in TBS containing 0.1% Tween-20 for 30 min and stained with primary and fluorescent dye-conjugated secondary antibodies diluted in the same solution for 1 h each, as detailed in the Key Resources Table. Immunoblots were imaged using an Odyssey CLx Imaging System (LI-COR).

### Blue native PAGE and immunoblot

Parasites were harvested as above, counted, and lysed in 750 mM aminocaproic acid, 0.5 mM EDTA, 50 mM Bis-Tris–HCl (pH 7.0), 2% (w/v) *n*-dodecyl-β-D-maltoside on ice for 10 min. Insoluble material was removed by centrifugation at 21,000 × *g* for 5 min at 4°C. The lysate was supplemented with Coomassie G-250 to 0.0625% and separated on a 4–16% gradient NativePAGE Bis-Tris gel (Invitrogen). Electrophoresis was performed at 80 V for 60 min using “dark” cathode buffer (1× NativePAGE Cathode Buffer Additive, 1× NativePAGE Running Buffer, both Invitrogen), followed by an additional 90 min at 250 V in “light” cathode buffer (0.1× NativePAGE Cathode Buffer Additive, 1× NativePAGE Running Buffer). Proteins were transferred to a charged 0.45 µm pore PVDF membrane (Merck) by wet transfer at 100 V for 1 h in Towbin buffer (25 mM Tris, 192 mM glycine, 10% methanol). Membranes were blocked with 5% (w/v) skim milk powder and stained with primary and horseradish peroxidase-conjugated secondary antibodies as detailed in the Key Resources Table and visualized using Pierce ECL Western Blotting Substrate (Thermo Scientific).

### Transmission electron microscopy

Intracellular parasites were fixed in 2.5% glutaraldehyde at 96 h after knockdown induction. Following serial washes in 0.1 M cacodylate buffer, samples were post-fixed in 1% OsO_4_ and 1.25% (v/v) potassium ferrocyanide in the same buffer for 1 h in the dark. Samples were washed with distilled water and contrasted en bloc with 0.5% aqueous uranyl acetate for 1 h at room temperature in the dark. Samples were dehydrated through an ascending acetone series (30%, 50%, 70%, 90%, 100%) and embedded in epoxy resin. Ultrathin sections (50 nm) were cut using a Leica ultramicrotome and collected on 100-mesh grids coated with Formvar. Sections were contrasted with 2% aqueous uranyl acetate and imaged using a JEOL 1400 Flash transmission electron microscope (JEOL, Japan) operated at 80 kV.

### TMRE assay

HFFs containing intracellular parasites were stained in 12.5 cm^2^ flasks with 1 mL of serum-free FluoroBrite medium (Gibco) containing 1 µM tetramethylrhodamine methyl ester (TMRE) and 3 µM Hoechst 33342 at 37°C for 30 min. In each replicate, one flask was treated simultaneously with 20 µM carbonyl cyanide m-chlorophenyl hydrazone (CCCP) as a positive control for mitochondrial depolarization. Working one flask at a time, parasites were mechanically released from host cells by syringe lysis, passed through a 5 µm filter to remove host cell debris, and immediately analyzed on a MACSQuant VYB flow cytometer (Miltenyi Biotec). Hoechst fluorescence was measured using 405 nm excitation with a 450/50 nm band-pass filter, and TMRE fluorescence using 561 nm excitation with a 615/20 nm band-pass filter. Data were processed in FlowJo v10.10.0. *Toxoplasma* cells were gated based on forward scatter, side scatter, singlets (FSC-H versus FSC-W), and Hoechst fluorescence. TMRE median fluorescence intensity (MFI) was quantified for gated parasites. Statistical differences between strains were determined using a two-sided paired *t*-test.

## DATA AVAILABILITY

Mass spectrometry proteomics data will be deposited to the ProteomeXchange Consortium via the PRIDE partner repository upon publication.

An interactive version of the protein–protein interaction network and associated predicted protein complex models is available at https://starpath.wi.mit.edu/.

All other data generated or analyzed in this study are included in the article and its supplementary data files. Plasmids and *Toxoplasma* strains generated in this study are available from the authors upon request.

## CODE AVAILABILITY

R scripts used to process the proteome-wide XL-MS data are available on GitHub (https://github.com/simonwbutterworth/Toxoplasma-XLMS-2026). Original Chai-1 model implementation and weights are available on GitHub (https://github.com/chaidiscovery/chai-lab). Modified Chai-1 inference script and Chai-1 prediction analysis scripts are available on GitHub (https://github.com/ashley-gin/starpath_toxo_ppi_network).

## ACKNOWLEDGMENTS

This work was supported by grants from the Gordon and Betty Moore Foundation (GBMF12274) and National Institutes of Health (AI197409) awarded to S.L., by a grant from the National Institutes of Health (AI193312) awarded to K.H., and a fellowship from the European Molecular Biology Organization (ALTF 312-2024) awarded to S.B.

## AUTHOR CONTRIBUTIONS

Conceptualization: S.B., S.L. Methodology: S.B., A.G., J.R. Investigation: S.B., A.G., S.S., I.T., J.R., T.D., V.S., L.L. Formal analysis: S.B., A.G. Visualization: S.B., A.G. Resources: F.S., K.H., L.S., S.O., S.L. Supervision: F.S., K.H., L.S., S.O., S.L. Funding acquisition: S.B., S.L. Writing – original draft: S.B., S.L. Writing – review & editing: all authors.

## SUPPLEMENTARY FIGURES

**Supplementary Figure 1.**
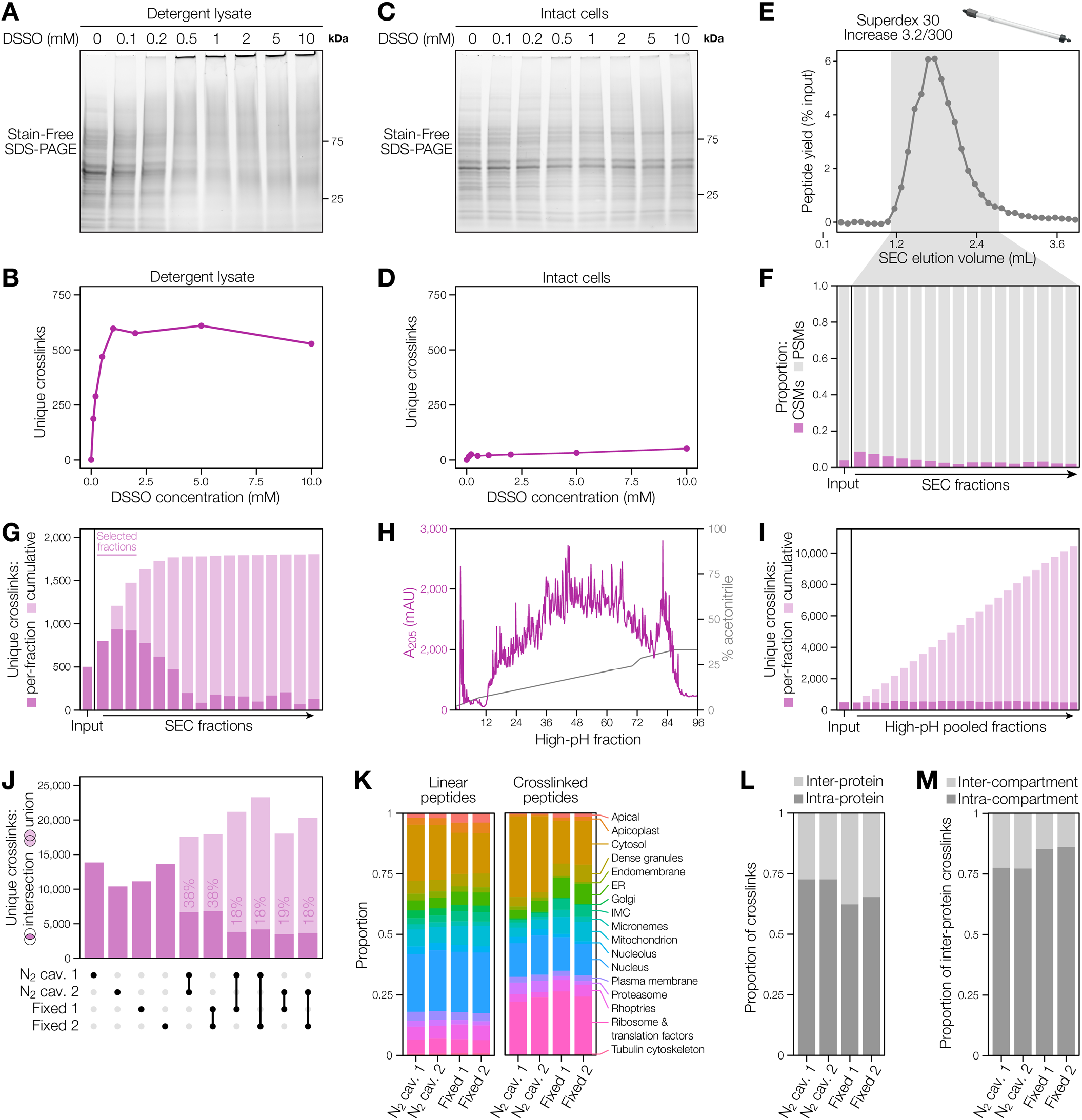
XL-MS method development. **(A)** SDS-PAGE of *Toxoplasma* detergent lysate crosslinked a range of DSSO concentrations. **(B)** Number of unique crosslinks identified from *Toxoplasma* detergent lysate crosslinked a range of DSSO concentrations. **(C)** SDS-PAGE of intact *Toxoplasma* cells crosslinked a range of DSSO concentrations. **(D)** Number of unique crosslinks identified from intact *Toxoplasma* cells crosslinked with a range of DSSO concentrations. **(E)** Representative elution profile from size-exclusion chromatography of crosslinked peptides. **(F)** Proportion of crosslink–spectrum matches (CSMs) versus peptide–spectrum matches (PSMs) for SEC fractions. **(G)** Unique and cumulative crosslinks identified across SEC fractions. **(H)** Representative ultraviolet absorbance chromatogram from high-pH reverse-phase HPLC fractiontion. **(I)** Unique and cumulative crosslinks identified across high-pH HPLC fraction pools. **(J)** Number of unique crosslinks identified in each XL-MS experiment and intersection and union of unique crosslinks for each experiment pair. Note that this is not an UpSet plot; each highlighted pair indicates the pair of datasets used to identify the union/intersection of unique crosslinks. **(K)** Subcellular distribution of linear and crosslinked peptides for each experimental replicate, based on hyperLOPIT spatial proteomics data supplemented with manual curation (see **Supplementary Data 3**). **(L)** Proportion of intra-versus inter-protein crosslinks for each replicate. **(M)** Proportion of inter-protein crosslinks linking proteins within the same subcellular compartment versus different compartments for each replicate.

**Supplementary Figure 2.**
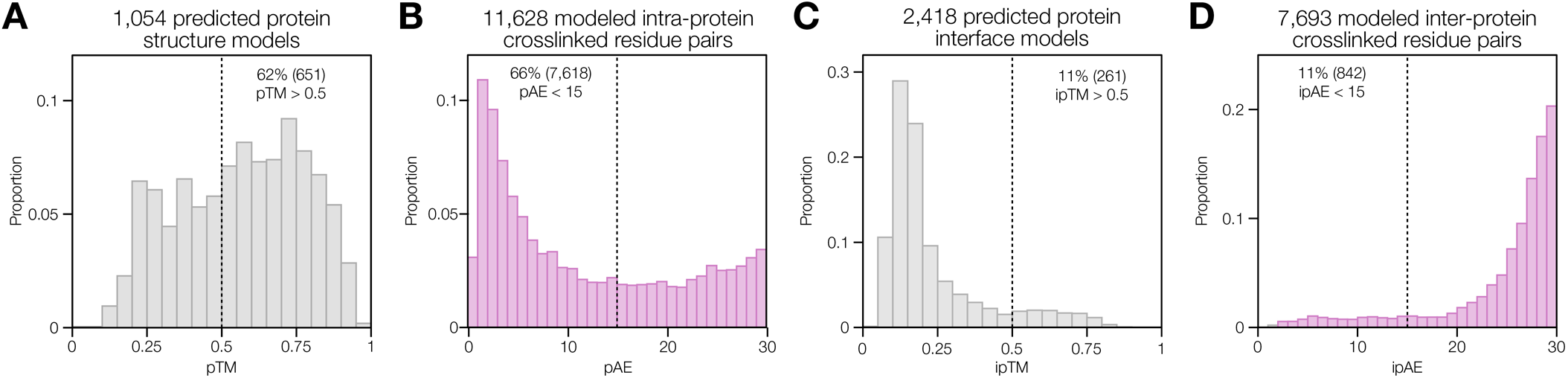
Protein structure modeling metrics. **(A)** Distribution of predicted template modeling (pTM) scores for Chai-1–predicted naive protein monomer structures. Scores greater than 0.5 indicate high confidence in the overall structure. **(B)** Distribution of mean predicted aligned error (pAE) scores for residue pairs identified by intra-protein crosslinks on predicted naive protein monomer structures. Mean pAE scores were calculated across the crosslinked residues and ±3 flanking residues. Scores less than 15 indicate high confidence in the positioning of residue pairs relative to one another. **(C)** Distribution of interface predicted template modeling (ipTM) scores for Chai-1–predicted naive protein heterodimer structures. Scores greater than 0.5 indicate high confidence in the relative positioning of the two protein subunits. **(D)** Distribution of mean interface predicted aligned error (ipAE) scores for residue pairs identified by inter-protein crosslinks on predicted naive heterodimers. Mean ipAE scores were calculated across the crosslinked residues and ±3 flanking residues. Scores less than 15 indicate high confidence in the positioning of residue pairs relative to one another.

**Supplementary Figure 3.**
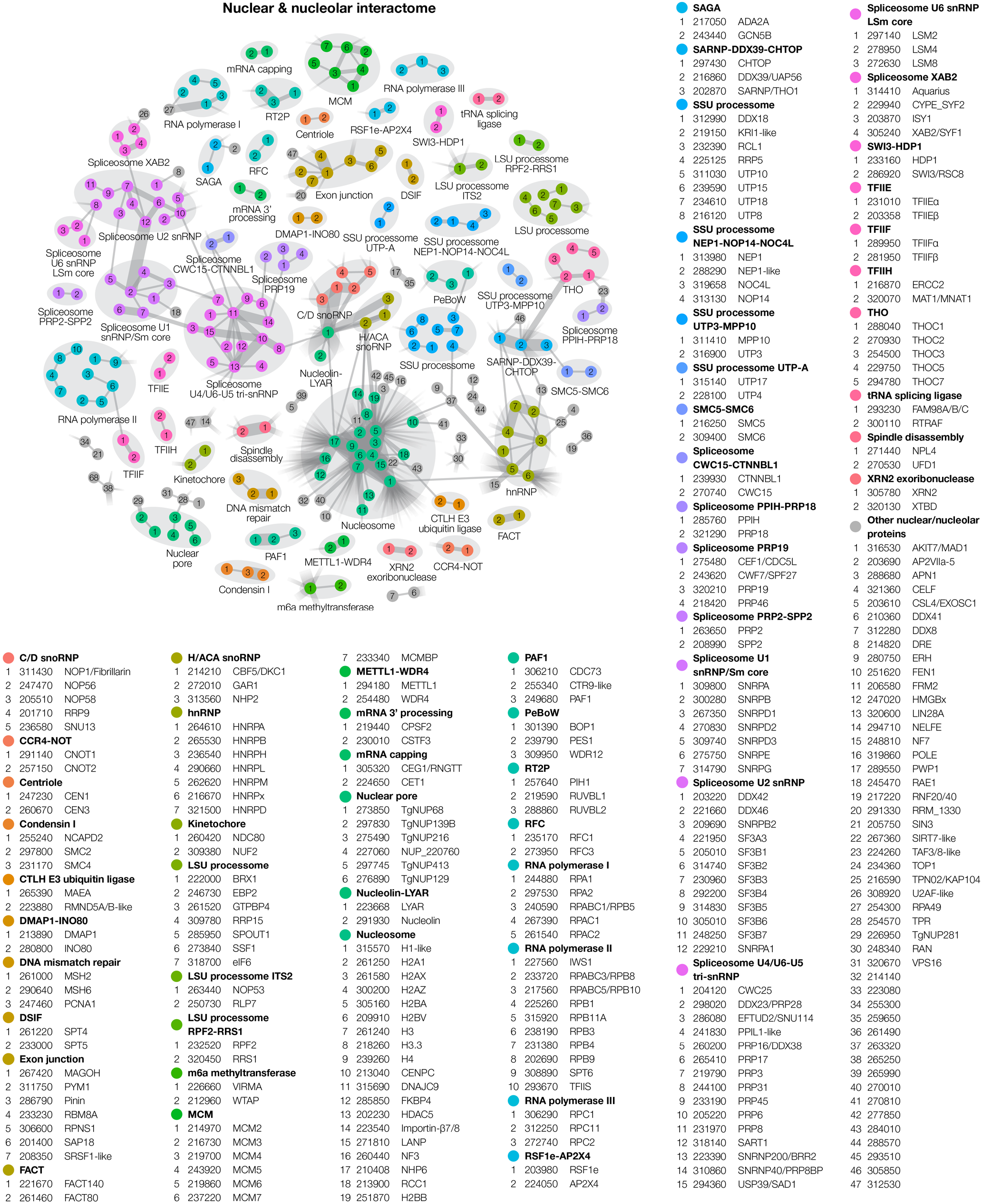
XL-MS-derived nuclear and nucleolar interactome.

**Supplementary Figure 4.**
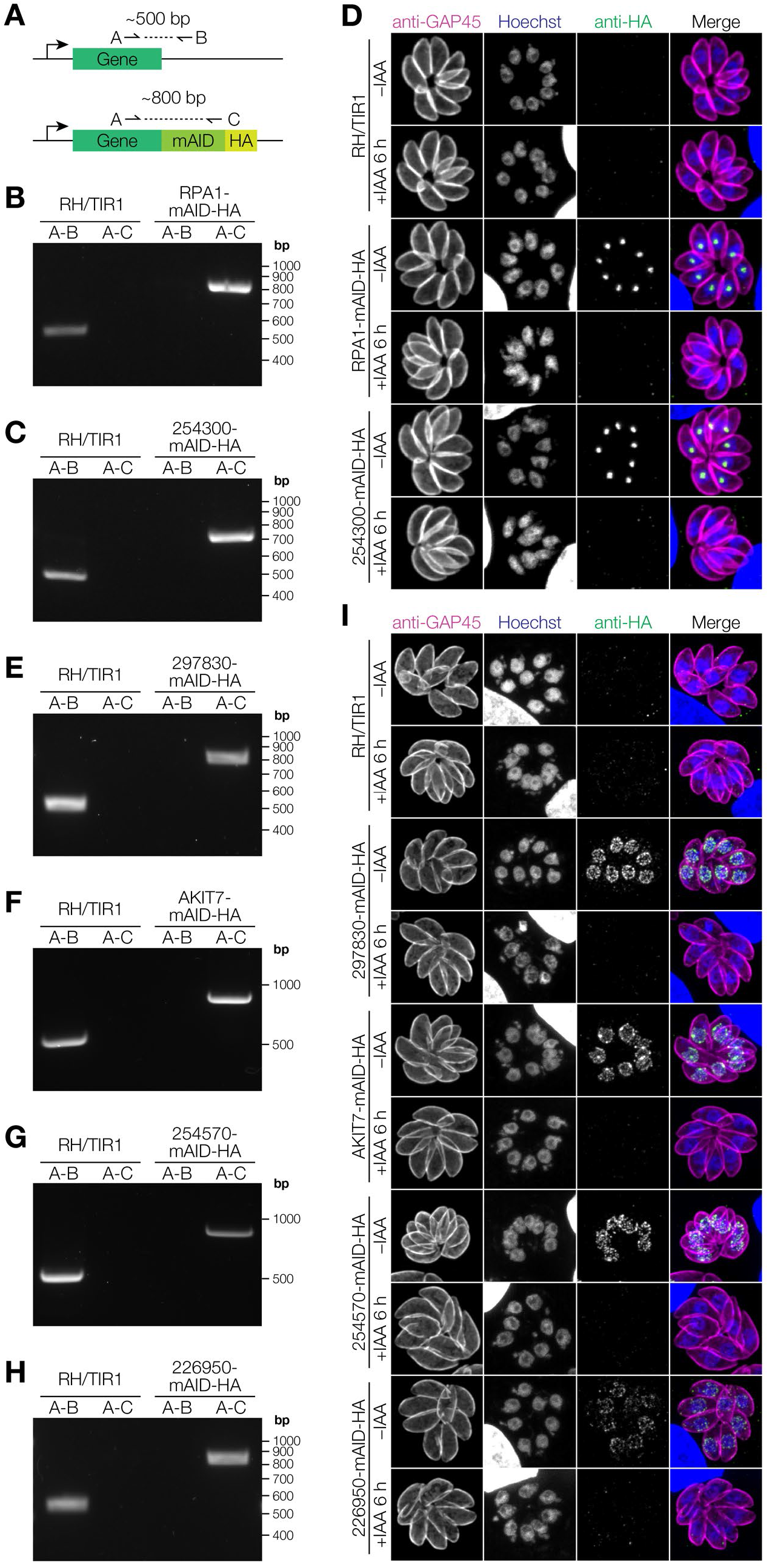
Validation of nuclear complex inducible knockdown strains. **(A)** PCR strategy to verify genomic integration of mAID-HA construct. **(B)** Diagnostic PCR to verify RPA1-mAID-HA strain. **(C)** Diagnostic PCR to verify 254300-mAID-HA strain. **(D)** Immunofluorescence microscopy validation of RPA1-mAID-HA and 254300-mAID-HA knockdown upon treatment with vehicle (–IAA) or auxin (+IAA) for 6 h. **(E)** Diagnostic PCR to verify 297830-mAID-HA strain. **(F)** Diagnostic PCR to verify AKIT7-mAID-HA strain. **(G)** Diagnostic PCR to verify 254570-mAID-HA strain. **(H)** Diagnostic PCR to verify 226950-mAID-HA strain. **(I)** Immunofluorescence microscopy validation of 297830-mAID-HA, AKIT7-mAID-HA, 254570-mAID-HA, and 226950-mAID-HA knockdown upon treatment with vehicle (– IAA) or auxin (+IAA) for 6 h.

**Supplementary Figure 5.**
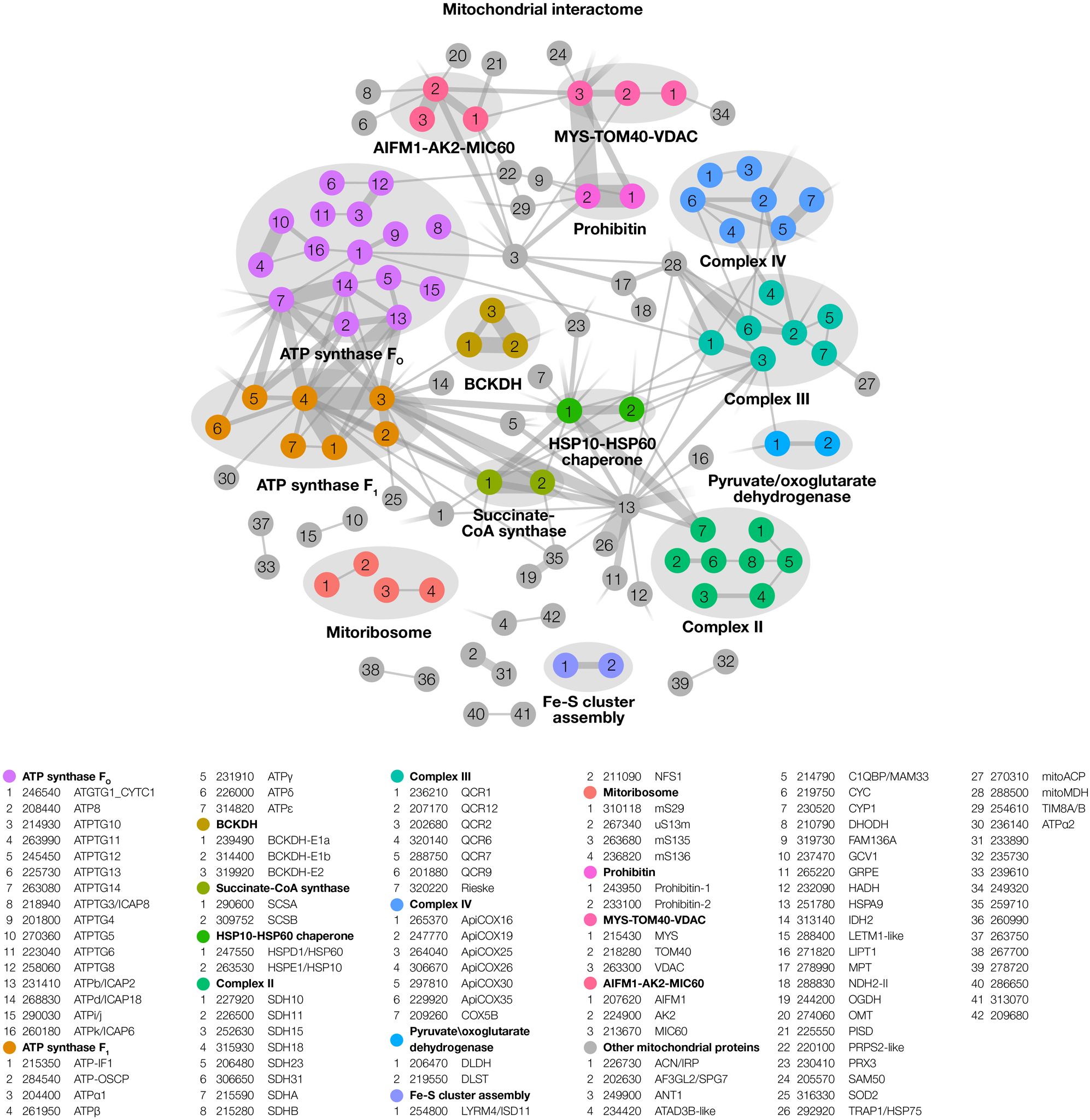
XL-MS-derived mitochondrial interactome.

**Supplementary Figure 6.**
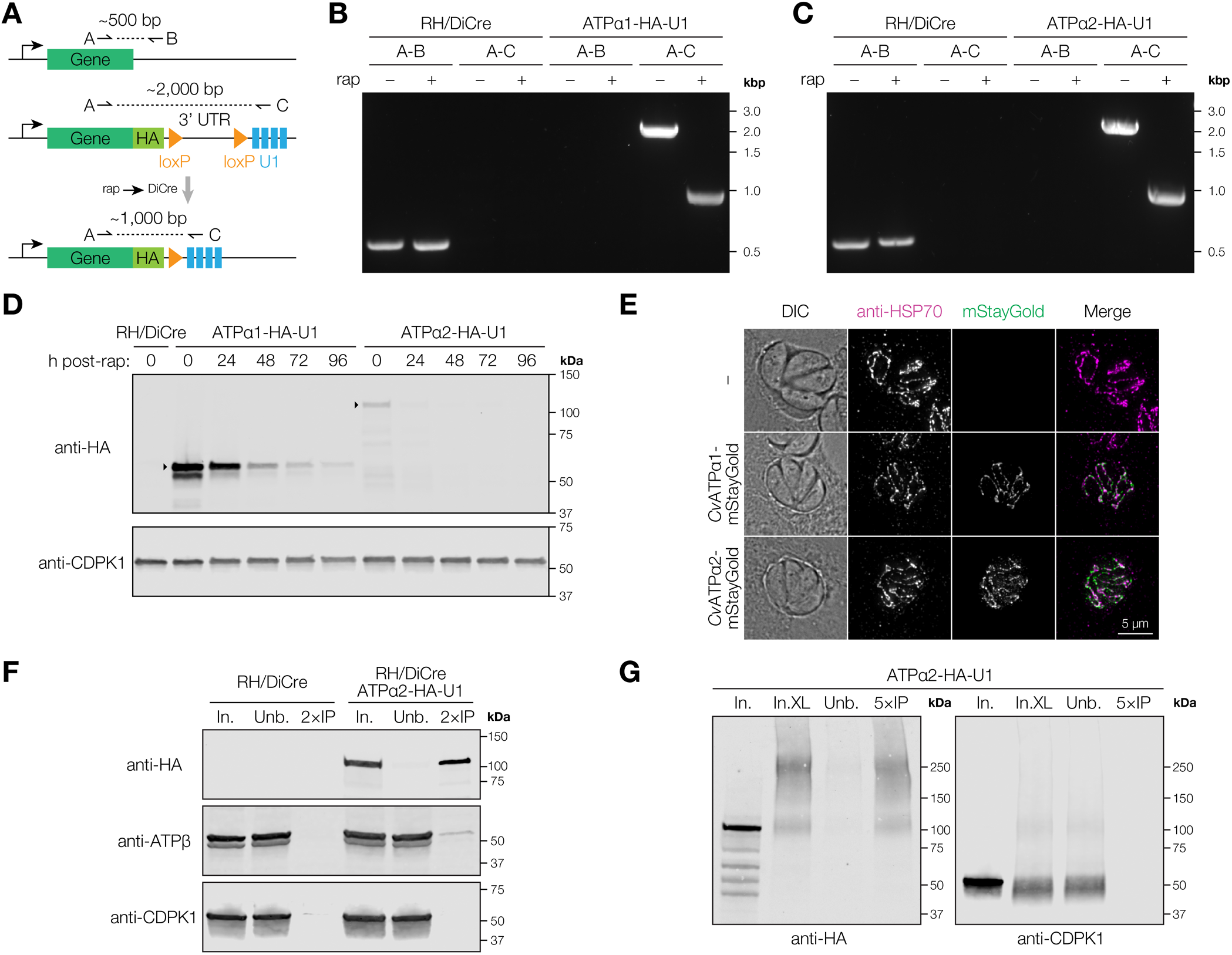
Validation of ATPα1 and ATPα2 inducible knockdown strains and *Chromera velia* expression constructs. **(A)** PCR strategy to verify genomic integration of HA-U1 construct and DiCre-mediated recombination. **(B)** Diagnostic PCR to verify ATPα1-HA-U1 strain. **(C)** Diagnostic PCR to verify ATPα2-HA-U1 strain. **(D)** Immunoblot analysis of ATPα1 and ATPα2 inducible knockdown kinetics. **(E)** Immunofluorescence microscopy of transiently expressed mStayGold-tagged *Chromera velia* ATPα1 and ATPα2 homologs in *Toxoplasma*. **(F)** Anti-HA co-immunoprecipitation and immunoblot of ATPα2. In. = input; Unb. = unbound; IP = immunoprecipitate. **(G)** Anti-HA immunoprecipitation of ATPα2 following DSSO crosslinking. In. = input prior to crosslinking; In. XL = DSSO crosslinked input; Unb. = unbound; IP = immunoprecipitate.

**Supplementary Figure 7.**
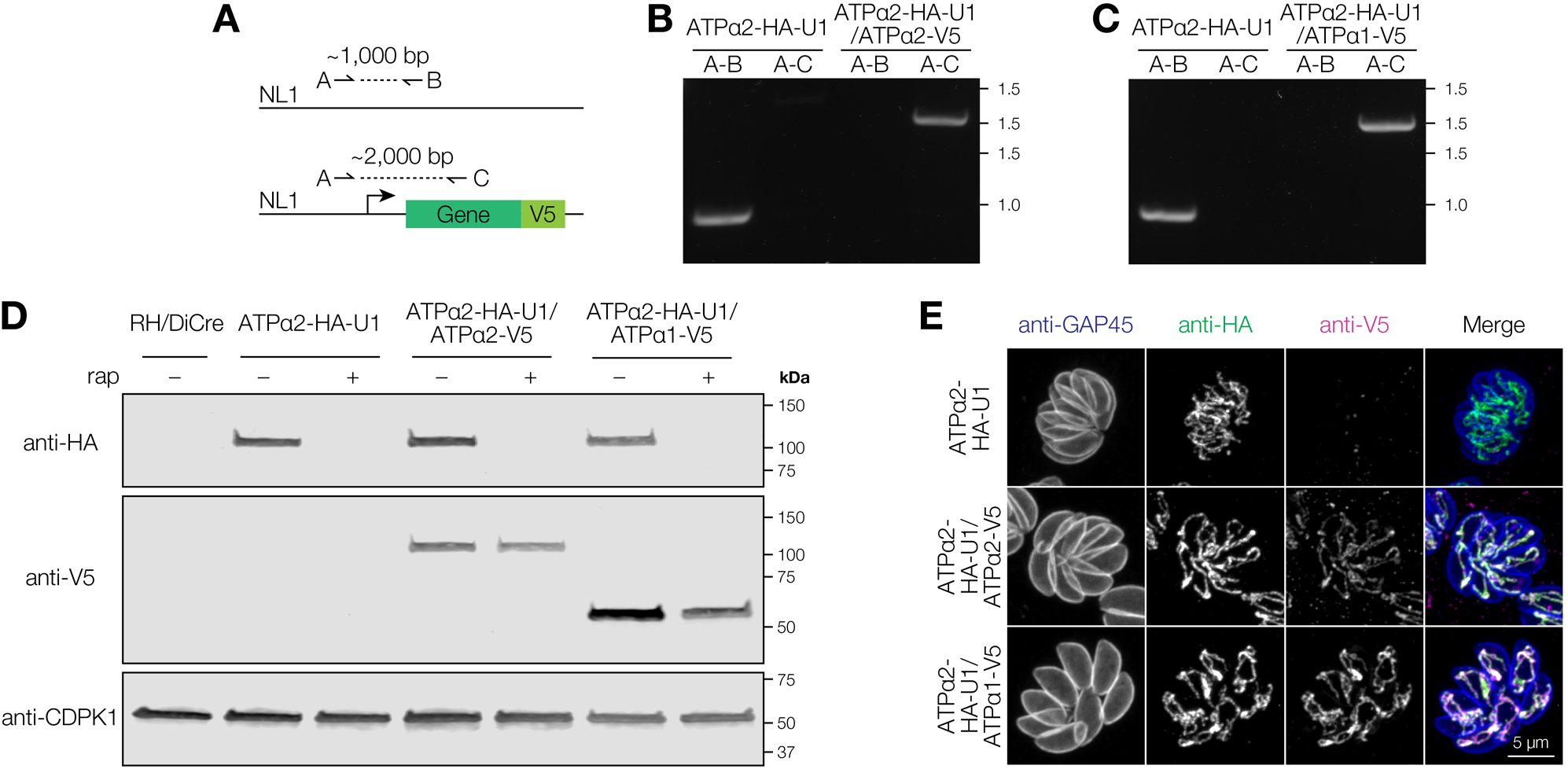
Validation of ATPα2-HA-U1 complementation strains. **(A)** PCR strategy to verify genomic integration of complementation constructs at Neutral Locus 1. **(B)** Diagnostic PCR to verify ATPα2-HA-U1/ATPα2-V5 strain. **(C)** Diagnostic PCR to verify ATPα2-HA-U1/ATPα1-V5 strain. **(D)** Immunoblot verifying tagged protein expression and rapamycin-induced knockdown in ATPα2-HA-U1/ATPα2-V5 and ATPα2-HA-U1/ATPα1-V5 strains. **(E)** Immunofluorescence microscopy demonstrated expected mitochondrial targeting of complemented V5-tagged proteins.

**Supplementary Figure 8.**
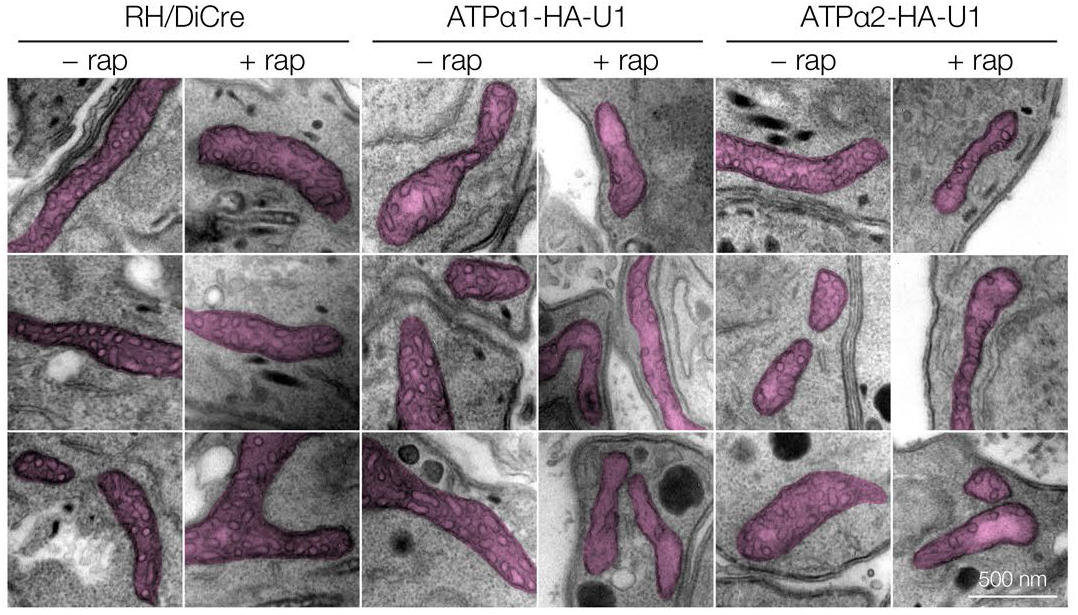
Transmission electron micrographs of the *Toxoplasma* mitochondrion following knockdown of ATPα1 and ATPα2.

## SUPPLEMENTARY DATA

**Supplementary Data 1. Combined XL-MS data from all experimental replicates**.

**(A)** Crosslink–spectrum matches identified at 1% FDR, including decoy matches. **(B)** Unique crosslinks, re-filtered to 1% FDR, including decoy matches. **(C)** Unique crosslinks following disambiguation of peptides mapping to multiple proteins as described in methods and re-indexing to the reference proteome. Remaining crosslinks with ambiguous peptides have been removed. **(D)** Unique residue pairs, re-filtered to 1% FDR. **(E)** Protein–protein interactions, re-filtered to 1% FDR. **(F)** Summary of score cutoffs and target/decoy counts.

**Supplementary Data 2. Chai-1 modeling results**.

**(A)** Intra-protein crosslink distances measured on predicted monomer structures. **(B)** Randomized lysine–lysine pair distances measured on predicted monomer structures. **(C)** Intra-protein crosslinks re-measured as inter-molecular links on predicted homodimer structures. **(D)** Inter-protein crosslink distances measured on naively predicted protein complex structures. **(E)** Randomized lysine–lysine pair distances measured on naively predicted protein complex structures. **(F)** Inter-protein crosslink distances measured on crosslink-constrained protein complex structures. Crosslinked residue pairs with mean ipAE > 15 in naive models were used to guide interface prediction specifying a maximum distance of 20 Å.

**Supplementary Data 3. Protein localization and complex annotation**.

**Supplementary Data 4. RPA1 and 254300 co-immunoprecipitation–mass spectrometry data**.

**(A)** DIA-NN output: precursors matrix. **(B)** DIA-NN output: protein groups matrix. **(C)** Protein-level processed data.

**Supplementary Data 5. α1 and α2 co-immunoprecipitation–mass spectrometry data**.

**(A)** DIA-NN output: precursors matrix. **(B)** DIA-NN output: protein groups matrix. **(C)** Protein-level processed data.

**Supplementary Data 6. α2 crosslinking–immunoprecipitation data**.

**(A)** Crosslink–spectrum matches identified at 1% FDR, including decoy matches. **(B)** Unique crosslinks, re-filtered to 1% FDR, including decoy matches. **(C)** Unique crosslinks between proteins of the α2 complex.

**Supplementary Data 7. Key Resources Table**.

## REFERENCES

1. Krogan, N.J., Cagney, G., Yu, H., Zhong, G., Guo, X., Ignatchenko, A., Li, J., Pu, S., Datta, N., Tikuisis, A.P., et al. (2006). Global landscape of protein complexes in the yeast Saccharomyces cerevisiae. Nature 440, 637–643.

2. Gavin, A.-C., Aloy, P., Grandi, P., Krause, R., Boesche, M., Marzioch, M., Rau, C., Jensen, L.J., Bastuck, S., Dümpelfeld, B., et al. (2006). Proteome survey reveals modularity of the yeast cell machinery. Nature 440, 631–636.

3. Cheng, F., Zhao, J., Wang, Y., Lu, W., Liu, Z., Zhou, Y., Martin, W.R., Wang, R., Huang, J., Hao, T., et al. (2021). Comprehensive characterization of protein–protein interactions perturbed by disease mutations. Nat. Genet. 53, 342–353.

4. Valastyan, J.S., and Lindquist, S. (2014). Mechanisms of protein-folding diseases at a glance. Dis. Model. Mech. 7, 9–14.

5. Shapira, S.D., Gat-Viks, I., Shum, B.O.V., Dricot, A., de Grace, M.M., Wu, L., Gupta, P.B., Hao, T., Silver, S.J., Root, D.E., et al. (2009). A physical and regulatory map of host-influenza interactions reveals pathways in H1N1 infection. Cell 139, 1255–1267.

6. Gordon, D.E., Jang, G.M., Bouhaddou, M., Xu, J., Obernier, K., White, K.M., O’Meara, M.J., Rezelj, V.V., Guo, J.Z., Swaney, D.L., et al. (2020). A SARS-CoV-2 protein interaction map reveals targets for drug repurposing. Nature 583, 459–468.

7. Greenblatt, J.F., Alberts, B.M., and Krogan, N.J. (2024). Discovery and significance of protein-protein interactions in health and disease. Cell 187, 6501–6517.

8. O’Reilly, F.J., and Rappsilber, J. (2018). Cross-linking mass spectrometry: methods and applications in structural, molecular and systems biology. Nat. Struct. Mol. Biol. 25, 1000–1008.

9. Wheat, A., Yu, C., Wang, X., Burke, A.M., Chemmama, I.E., Kaake, R.M., Baker, P., Rychnovsky, S.D., Yang, J., and Huang, L. (2021). Protein interaction landscapes revealed by advanced in vivo cross-linking-mass spectrometry. Proc. Natl. Acad. Sci. U. S. A. 118, e2023360118.

10. Bartolec, T.K., Vázquez-Campos, X., Norman, A., Luong, C., Johnson, M., Payne, R.J., Wilkins, M.R., Mackay, J.P., and Low, J.K.K. (2023). Cross-linking mass spectrometry discovers, evaluates, and corroborates structures and protein–protein interactions in the human cell. Proc. Natl. Acad. Sci. U. S. A. 120, e2219418120.

11. Gonzalez-Lozano, M.A., Schmid, E.W., Miguel Whelan, E., Jiang, Y., Paulo, J.A., Walter, J.C., and Harper, J.W. (2025). EndoMAP.v1 charts the structural landscape of human early endosome complexes. Nature 643, 252–261.

12. Pappas, G., Roussos, N., and Falagas, M.E. (2009). Toxoplasmosis snapshots: global status of Toxoplasma gondii seroprevalence and implications for pregnancy and congenital toxoplasmosis. Int. J. Parasitol. 39, 1385–1394.

13. Montoya, J.G., and Liesenfeld, O. (2004). Toxoplasmosis. Lancet 363, 1965–1976.

14. Escalante, A.A., and Ayala, F.J. (1995). Evolutionary origin of Plasmodium and other Apicomplexa based on rRNA genes. Proc. Natl. Acad. Sci. U. S. A. 92, 5793–5797.

15. Kim, K., and Weiss, L.M. (2004). Toxoplasma gondii: the model apicomplexan. Int. J. Parasitol. 34, 423–432.

16. Tonkin, M.L., Arredondo, S.A., Loveless, B.C., Serpa, J.J., Makepeace, K.A.T., Sundar, N., Petrotchenko, E.V., Miller, L.H., Grigg, M.E., and Boulanger, M.J. (2013). Structural and biochemical characterization of Plasmodium falciparum 12 (Pf12) reveals a unique interdomain organization and the potential for an antiparallel arrangement with Pf41. J. Biol. Chem. 288, 12805–12817.

17. Wong, W., Webb, A.I., Olshina, M.A., Infusini, G., Tan, Y.H., Hanssen, E., Catimel, B., Suarez, C., Condron, M., Angrisano, F., et al. (2014). A mechanism for actin filament severing by malaria parasite actin depolymerizing factor 1 via a low affinity binding interface. J. Biol. Chem. 289, 4043–4054.

18. Jagadeeshaprasad, M.G., Bewley, M.C., Lovely, G., Flanagan, J.M., and Gowda, C.D. (2022). Chemical crosslinking and peptide mapping reveal structural characteristics of interdomain 1 and 2 of the placental malaria adherent protein VAR2CSA. FASEB J. 36 Suppl 1. 10.1096/fasebj.2022.36.S1.R2112.

19. Tomita, T., Weyer, E., Guevara, R.B., Sidoli, S., Aguilan, J.T., and Weiss, L.M. (2025). Mapping a Toxoplasma gondii interactome by crosslinking mass spectrometry and machine learning. mBio 16, e0215925.

20. Mostafavi, S., Maurer, V., Ruwolt, M., Schwartz, J., Stein, F., Niedermüller, K., Gilberger, T.W., Liu, F., Seuring, C., Witt, S., et al. (2026). Deciphering membrane protein complexes in Plasmodium falciparum gametocytes via integrative structural systems biology. Mol. Cell. Proteomics 25, 101567.

21. Kao, A., Chiu, C.-L., Vellucci, D., Yang, Y., Patel, V.R., Guan, S., Randall, A., Baldi, P., Rychnovsky, S.D., and Huang, L. (2011). Development of a novel cross-linking strategy for fast and accurate identification of cross-linked peptides of protein complexes. Mol. Cell. Proteomics 10, M110.002212.

22. Simpson, R.J. (2010). Disruption of cultured cells by nitrogen cavitation. Cold Spring Harb. Protoc. 2010, db.prot5513.

23. Michael, A.R.M., Amaral, B.C., Ball, K.L., Eiriksson, K.H., and Schriemer, D.C. (2024). Cell fixation improves performance of in situ crosslinking mass spectrometry while preserving cellular ultrastructure. Nat. Commun. 15, 8537.

24. Leitner, A., Reischl, R., Walzthoeni, T., Herzog, F., Bohn, S., Förster, F., and Aebersold, R. (2012). Expanding the chemical cross-linking toolbox by the use of multiple proteases and enrichment by size exclusion chromatography. Mol. Cell. Proteomics 11, M111.014126.

25. Stieger, C.E., Doppler, P., and Mechtler, K. (2019). Optimized fragmentation improves the identification of peptides cross-linked by MS-cleavable reagents. J. Proteome Res. 18, 1363–1370.

26. Mühleip, A., Kock Flygaard, R., Ovciarikova, J., Lacombe, A., Fernandes, P., Sheiner, L., and Amunts, A. (2021). ATP synthase hexamer assemblies shape cristae of Toxoplasma mitochondria. Nat. Commun. 12, 120.

27. Merkley, E.D., Rysavy, S., Kahraman, A., Hafen, R.P., Daggett, V., and Adkins, J.N. (2014). Distance restraints from crosslinking mass spectrometry: mining a molecular dynamics simulation database to evaluate lysine-lysine distances. Protein Sci. 23, 747–759.

28. MacLean, A.E., Shikha, S., Ferreira Silva, M., Gramelspacher, M.J., Nilsen, A., Liebman, K.M., Pou, S., Winter, R.W., Meir, A., Riscoe, M.K., et al. (2025). Structure, assembly and inhibition of the Toxoplasma gondii respiratory chain supercomplex. Nat. Struct. Mol. Biol. 32, 1424–1433.

29. Wernimont, A.K., Artz, J.D., Finerty, P., Jr, Lin, Y.-H., Amani, M., Allali-Hassani, A., Senisterra, G., Vedadi, M., Tempel, W., Mackenzie, F., et al. (2010). Structures of apicomplexan calcium-dependent protein kinases reveal mechanism of activation by calcium. Nat. Struct. Mol. Biol. 17, 596–601.

30. Chen, Z.A., and Rappsilber, J. (2023). Protein structure dynamics by crosslinking mass spectrometry. Curr. Opin. Struct. Biol. 80, 102599.

31. Chai Discovery, Boitreaud, J., Dent, J., McPartlon, M., Meier, J., Reis, V., Rogozhnikov, A., and Wu, K. (2024). Chai-1: Decoding the molecular interactions of life. bioRxiv. 10.1101/2024.10.10.615955.

32. Schweke, H., Pacesa, M., Levin, T., Goverde, C.A., Kumar, P., Duhoo, Y., Dornfeld, L.J., Dubreuil, B., Georgeon, S., Ovchinnikov, S., et al. (2024). An atlas of protein homo-oligomerization across domains of life. Cell 187, 999–1010.e15.

33. Forshaw, T.E., Reisz, J.A., Nelson, K.J., Gumpena, R., Lawson, J.R., Jönsson, T.J., Wu, H., Clodfelter, J.E., Johnson, L.C., Furdui, C.M., et al. (2021). Specificity of human sulfiredoxin for reductant and peroxiredoxin oligomeric state. Antioxidants (Basel) 10, 946.

34. Tomko, R.J., Jr, Funakoshi, M., Schneider, K., Wang, J., and Hochstrasser, M. (2010). Heterohexameric ring arrangement of the eukaryotic proteasomal ATPases: implications for proteasome structure and assembly. Mol. Cell 38, 393–403.

35. Schweitzer, A., Aufderheide, A., Rudack, T., Beck, F., Pfeifer, G., Plitzko, J.M., Sakata, E., Schulten, K., Förster, F., and Baumeister, W. (2016). Structure of the human 26S proteasome at a resolution of 3.9 Å. Proc. Natl. Acad. Sci. U. S. A. 113, 7816–7821.

36. Williams, M.J., Alonso, H., Enciso, M., Egarter, S., Sheiner, L., Meissner, M., Striepen, B., Smith, B.J., and Tonkin, C.J. (2015). Two essential light chains regulate the MyoA lever arm to promote Toxoplasma gliding motility. mBio 6, e00845–15.

37. Yook, S.-H., Oltvai, Z.N., and Barabási, A.-L. (2004). Functional and topological characterization of protein interaction networks. Proteomics 4, 928–942.

38. Clauset, A., Shalizi, C.R., and Newman, M.E.J. (2007). Power-law distributions in empirical data. arXiv:0706.1062.

39. Jeong, H., Mason, S.P., Barabási, A.L., and Oltvai, Z.N. (2001). Lethality and centrality in protein networks. Nature 411, 41–42.

40. Barylyuk, K., Koreny, L., Ke, H., Butterworth, S., Crook, O.M., Lassadi, I., Gupta, V., Tromer, E., Mourier, T., Stevens, T.J., et al. (2020). A comprehensive subcellular atlas of the Toxoplasma proteome via hyperLOPIT provides spatial context for protein functions. Cell Host Microbe 28, 752–766.e9.

41. Goldfarb, D.S., Corbett, A.H., Mason, D.A., Harreman, M.T., and Adam, S.A. (2004). Importin alpha: a multipurpose nuclear-transport receptor. Trends Cell Biol. 14, 505–514.

42. Cao, K., Nakajima, R., Meyer, H.H., and Zheng, Y. (2003). The AAA-ATPase Cdc48/p97 regulates spindle disassembly at the end of mitosis. Cell 115, 355–367.

43. Ye, Y., Meyer, H.H., and Rapoport, T.A. (2001). The AAA ATPase Cdc48/p97 and its partners transport proteins from the ER into the cytosol. Nature 414, 652–656.

44. Nishimura, K., Fukagawa, T., Takisawa, H., Kakimoto, T., and Kanemaki, M. (2009). An auxin-based degron system for the rapid depletion of proteins in nonplant cells. Nat. Methods 6, 917–922.

45. Sidik, S.M., Huet, D., Ganesan, S.M., Huynh, M.-H., Wang, T., Nasamu, A.S., Thiru, P., Saeij, J.P.J., Carruthers, V.B., Niles, J.C., et al. (2016). A genome-wide CRISPR screen in Toxoplasma identifies essential apicomplexan genes. Cell 166, 1423–1435.e12.

46. Carré, C., and Shiekhattar, R. (2011). Human GTPases associate with RNA polymerase II to mediate its nuclear import. Mol. Cell. Biol. 31, 3953–3962.

47. Zimmermann, L., Stephens, A., Nam, S.-Z., Rau, D., Kübler, J., Lozajic, M., Gabler, F., Söding, J., Lupas, A.N., and Alva, V. (2018). A completely reimplemented MPI bioinformatics toolkit with a new HHpred server at its core. J. Mol. Biol. 430, 2237–2243.

48. Misiaszek, A.D., Girbig, M., Grötsch, H., Baudin, F., Murciano, B., Lafita, A., and Müller, C.W. (2021). Cryo-EM structures of human RNA polymerase I. Nat. Struct. Mol. Biol. 28, 997–1008.

49. Singh, D., Soni, N., Hutchings, J., Echeverria, I., Shaikh, F., Duquette, M., Suslov, S., Li, Z., van Eeuwen, T., Molloy, K., et al. (2024). The molecular architecture of the nuclear basket. Cell 187, 5267–5281.e13.

50. Dewangan, P.S., Dohr, S.R., Trotter, J., and Reese, M.L. (2025). Identification of divergent Toxoplasma nuclear pore complex components highlights speciation of mRNA export machinery. bioRxiv. 10.1101/2025.08.27.672535.

51. Brusini, L., Dos Santos Pacheco, N., Tromer, E.C., Soldati-Favre, D., and Brochet, M. (2022). Composition and organization of kinetochores show plasticity in apicomplexan chromosome segregation. J. Cell Biol. 221, e202111084.

52. Campbell, M.S., Chan, G.K., and Yen, T.J. (2001). Mitotic checkpoint proteins HsMAD1 and HsMAD2 are associated with nuclear pore complexes in interphase. J. Cell Sci. 114, 953–963.

53. Iouk, T., Kerscher, O., Scott, R.J., Basrai, M.A., and Wozniak, R.W. (2002). The yeast nuclear pore complex functionally interacts with components of the spindle assembly checkpoint. J. Cell Biol. 159, 807–819.

54. Hutten, S., and Kehlenbach, R.H. (2006). Nup214 is required for CRM1-dependent nuclear protein export in vivo. Mol. Cell. Biol. 26, 6772–6785.

55. Gorsch, L.C., Dockendorff, T.C., and Cole, C.N. (1995). A conditional allele of the novel repeat-containing yeast nucleoporin RAT7/NUP159 causes both rapid cessation of mRNA export and reversible clustering of nuclear pore complexes. J. Cell Biol. 129, 939–955.

56. Maclean, A.E., Bridges, H.R., Silva, M.F., Ding, S., Ovciarikova, J., Hirst, J., and Sheiner, L. (2021). Complexome profile of Toxoplasma gondii mitochondria identifies divergent subunits of respiratory chain complexes including new subunits of cytochrome bc1 complex. PLoS Pathog. 17, e1009301.

57. Huet, D., Rajendran, E., van Dooren, G.G., and Lourido, S. (2018). Identification of cryptic subunits from an apicomplexan ATP synthase. eLife 7, e38097.

58. Salunke, R., Mourier, T., Banerjee, M., Pain, A., and Shanmugam, D. (2018). Highly diverged novel subunit composition of apicomplexan F-type ATP synthase identified from Toxoplasma gondii. PLoS Biol. 16, e2006128.

59. Pieperhoff, M.S., Pall, G.S., Jiménez-Ruiz, E., Das, S., Melatti, C., Gow, M., Wong, E.H., Heng, J., Müller, S., Blackman, M.J., et al. (2015). Conditional U1 gene silencing in Toxoplasma gondii. PLoS One 10, e0130356.

60. Tengganu, I.F., and Hu, K. (2026). Transfection of the free-living alga Chromera velia enables direct comparisons with its parasitic apicomplexan relative, Toxoplasma gondii. J. Cell Sci. 139, jcs264400.

61. Markus, B.M., Bell, G.W., Lorenzi, H.A., and Lourido, S. (2019). Optimizing systems for Cas9 expression in Toxoplasma gondii. mSphere 4, e00386–19.

62. Crowley, L.C., Christensen, M.E., and Waterhouse, N.J. (2016). Measuring mitochondrial transmembrane potential by TMRE staining. Cold Spring Harb. Protoc. 2016, db.prot087361.

63. Usey, M.M., and Huet, D. (2023). ATP synthase-associated coiled-coil-helix-coiled-coil-helix (CHCH) domain-containing proteins are critical for mitochondrial function in Toxoplasma gondii. mBio 14, e0176923.

64. Lenz, S., Sinn, L.R., O’Reilly, F.J., Fischer, L., Wegner, F., and Rappsilber, J. (2021). Reliable identification of protein-protein interactions by crosslinking mass spectrometry. Nat. Commun. 12, 3564.

65. Huttlin, E.L., Bruckner, R.J., Navarrete-Perea, J., Cannon, J.R., Baltier, K., Gebreab, F., Gygi, M.P., Thornock, A., Zarraga, G., Tam, S., et al. (2021). Dual proteome-scale networks reveal cell-specific remodeling of the human interactome. Cell 184, 3022–3040.e28.

66. Havugimana, P.C., Hart, G.T., Nepusz, T., Yang, H., Turinsky, A.L., Li, Z., Wang, P.I., Boutz, D.R., Fong, V., Phanse, S., et al. (2012). A census of human soluble protein complexes. Cell 150, 1068–1081.

67. Swapna, L.S., Stevens, G.C., Sardinha-Silva, A., Hu, L.Z., Brand, V., Fusca, D.D., Wan, C., Xiong, X., Boyle, J.P., Grigg, M.E., et al. (2024). ToxoNet: A high confidence map of protein-protein interactions in Toxoplasma gondii. PLoS Comput. Biol. 20, e1012208.

68. Han, Y., Tsenkov, M.I., Venanzi, N.A.E., Bertoni, D., Cha, S., Chacón, A., Dietrich, N., Fomitchev, B., Goldtzvik, Y., Hsu, D., et al. (2026). AlphaFold Database expands to proteome-scale quaternary structures. bioRxiv. 10.64898/2026.03.27.714458.

69. Abramson, J., Adler, J., Dunger, J., Evans, R., Green, T., Pritzel, A., Ronneberger, O., Willmore, L., Ballard, A.J., Bambrick, J., et al. (2024). Accurate structure prediction of biomolecular interactions with AlphaFold 3. Nature 630, 493–500.

70. Stahl, K., Graziadei, A., Dau, T., Brock, O., and Rappsilber, J. (2023). Protein structure prediction with in-cell photo-crosslinking mass spectrometry and deep learning. Nat. Biotechnol. 41, 1810–1819.

71. Gilep, K., Obarska-Kosinska, A., and Kosinski, J. (2024). Improving AlphaFold 3 structural modeling by incorporating explicit crosslinks. bioRxiv. 10.1101/2024.12.03.626671.

72. Mair, G., Daiß, J.L., Engel, C., and Panov, K.I. (2025). The ribosomal RNA transcription landscapes of Plasmodium falciparum and related apicomplexan parasites. Nucleic Acids Res. 53, gkaf641.

73. Courjol, F., Mouveaux, T., Lesage, K., Saliou, J.-M., Werkmeister, E., Bonabaud, M., Rohmer, M., Slomianny, C., Lafont, F., and Gissot, M. (2017). Characterization of a nuclear pore protein sheds light on the roles and composition of the Toxoplasma gondii nuclear pore complex. Cell. Mol. Life Sci. 74, 2107–2125.

74. Ovciarikova, J., Shikha, S., Lacombe, A., Courjol, F., McCrone, R., Hussain, W., Maclean, A., Lemgruber, L., Martins-Duarte, E.S., Gissot, M., et al. (2024). Two ancient membrane pores mediate mitochondrial-nucleus membrane contact sites. J. Cell Biol. 223, e202304075.

75. Onischenko, E., Noor, E., Fischer, J.S., Gillet, L., Wojtynek, M., Vallotton, P., and Weis, K. (2020). Maturation kinetics of a multiprotein complex revealed by metabolic labeling. Cell 183, 1785–1800.e26.

76. Hurwitz, M.E., Strambio-de-Castillia, C., and Blobel, G. (1998). Two yeast nuclear pore complex proteins involved in mRNA export form a cytoplasmically oriented subcomplex. Proc. Natl. Acad. Sci. U. S. A. 95, 11241–11245.

77. Rühle, T., and Leister, D. (2015). Assembly of F1F0-ATP synthases. Biochim. Biophys. Acta 1847, 849–860.

78. Walker, J.E. (2013). The ATP synthase: the understood, the uncertain and the unknown. Biochem. Soc. Trans. 41, 1–16.

79. Ruwolt, M., Schnirch, L., Borges Lima, D., Nadler-Holly, M., Viner, R., and Liu, F. (2022). Optimized TMT-based quantitative cross-linking mass spectrometry strategy for large-scale interactomic studies. Anal. Chem. 94, 5265–5272.

80. Hughes, C.S., Moggridge, S., Müller, T., Sorensen, P.H., Morin, G.B., and Krijgsveld, J. (2019). Single-pot, solid-phase-enhanced sample preparation for proteomics experiments. Nat. Protoc. 14, 68–85.

81. Liu, F., Rijkers, D.T.S., Post, H., and Heck, A.J.R. (2015). Proteome-wide profiling of protein assemblies by cross-linking mass spectrometry. Nat. Methods 12, 1179–1184.

82. Liu, F., Lössl, P., Scheltema, R., Viner, R., and Heck, A.J.R. (2017). Optimized fragmentation schemes and data analysis strategies for proteome-wide cross-link identification. Nat. Commun. 8, 15473.

83. Smith, T.A., Lopez-Perez, G.S., Herneisen, A.L., Shortt, E., and Lourido, S. (2022). Screening the Toxoplasma kinome with high-throughput tagging identifies a regulator of invasion and egress. Nat. Microbiol. 7, 868–881.

84. Hunt, A., Russell, M.R.G., Wagener, J., Kent, R., Carmeille, R., Peddie, C.J., Collinson, L., Heaslip, A., Ward, G.E., and Treeck, M. (2019). Differential requirements for cyclase-associated protein (CAP) in actin-dependent processes of Toxoplasma gondii. eLife 8, e50598.

85. Kumar, S., Stecher, G., Suleski, M., Sanderford, M., Sharma, S., and Tamura, K. (2024). MEGA12: Molecular evolutionary genetic analysis version 12 for adaptive and green computing. Mol. Biol. Evol. 41, msae263.

86. Schindelin, J., Arganda-Carreras, I., Frise, E., Kaynig, V., Longair, M., Pietzsch, T., Preibisch, S., Rueden, C., Saalfeld, S., Schmid, B., et al. (2012). Fiji: an open-source platform for biological-image analysis. Nat. Methods 9, 676–682.

87. HaileMariam, M., Eguez, R.V., Singh, H., Bekele, S., Ameni, G., Pieper, R., and Yu, Y. (2018). S-Trap, an ultrafast sample-preparation approach for shotgun proteomics. J. Proteome Res. 17, 2917–2924.

88. Alvarez-Jarreta, J., Amos, B., Aurrecoechea, C., Bah, S., Barba, M., Barreto, A., Basenko, E.Y., Belnap, R., Blevins, A., Böhme, U., et al. (2024). VEuPathDB: the eukaryotic pathogen, vector and host bioinformatics resource center in 2023. Nucleic Acids Res. 52, D808–D816.

89. Li, Z., Guo, Q., Zheng, L., Ji, Y., Xie, Y.-T., Lai, D.-H., Lun, Z.-R., Suo, X., and Gao, N. (2017). Cryo-EM structures of the 80S ribosomes from human parasites Trichomonas vaginalis and Toxoplasma gondii. Cell Res. 27, 1275–1288.

